# Organ-specific isolation of hepatocyte extracellular vesicles from human plasma enables tissue-resolved proteomic and miRNA profiling

**DOI:** 10.1101/2025.06.13.658908

**Authors:** Ricardo Figueiras, Susana Vagueiro, Raissa Kay, Elnaz Persia, Roger de Alwis, Razan Hijazi, Brian Davidson, Raquel Sanches-Kuiper, Pierre Arsène, Tomás Dias

**Affiliations:** Mursla Bio, Cambridge, UK & Cambridge, US; University College London, UK

**Keywords:** Extracellular Vesicles (EVs), Liquid Biopsy, Precision Medicine, Hepatocellular Carcinoma (HCC), Organ-Specific EVs, Multi-Omics, EV Proteomics, miRNA Biomarkers, Immunocapture, NEXPLOR Platform

## Abstract

Tissue-specific resolution remains a key limitation in liquid biopsy to achieve the highest accuracy for precision medicine. To address this limitation, we developed NEXPLOR (Novel EXtracellular vesicle PopuLation and Omics Revealer), a magnetic bead-based platform enabling selective immuno-isolation of tissue-derived EVs. Focusing on hepatocyte-derived EVs (h-EVs), we demonstrate NEXPLOR’s specificity and sensitivity in capturing rare EV subpopulations directly from human plasma. Using *in silico* tissue marker discovery, we identified and validated a TOP4 capture antibody panel (ASGR1, ASGR2, TFR2, SLCO1B1) for h-EV isolation through an ultrasensitive orthogonal method (O-NEXOS). Applied to liver disease using clinical plasma samples, NEXPLOR enabled deeper and more reproducible proteomic profiling compared to matched bulk EVs, revealing liver-specific and disease-relevant pathways, including ferroptosis, HIF-1α signaling, and central carbon metabolism in cancer. Moreover, small RNA sequencing uncovered a reduced but highly informative set of miRNAs, including *miR-124-3p* and *miR-23b-3p*, differentially expressed in hepatocellular carcinoma (HCC) compared to cirrhosis, and undetectable in bulk EVs. These molecular signatures suggest disease state based on real-time hepatocyte biology. Our findings establish NEXPLOR as a robust platform for tissue-specific EV capture, enabling tissue-resolved, multi-omic biomarker discovery. This opens new frontiers in early disease detection, longitudinal monitoring, and AI-powered biomarker discovery for precision medicine.

## INTRODUCTION

Precision medicine increasingly depends on liquid biopsy for non-invasive, repeatable access to molecular information from blood. Yet restoring tissue specificity and the associated spatial resolution remains a critical challenge. A promising path to address this challenge lies in circulating extracellular vesicles (EVs) and other extracellular particles (EPs), which offer a uniquely convenient and powerful window into the physiological and pathological state of their cells of origin, including those from tissues and organs ^1, 2, 3–5^.

EVs and other EPs carry proteins, lipids, metabolites, and nucleic acids. Unlike cell-free and circulating tumor DNA (cf/ctDNA), which is primarily released through apoptosis or necrosis, typically at intervals ranging from days to months depending on the cell type, EVs and other EPs are secreted continuously, providing a multi-omic snapshot of the living cellular *milieu*, with more than 1,000 EVs released per cell per day ^6,7^.

Additionally, EVs contain tissue-specific markers, enabling the molecular cargo they carry to be linked to their tissue of origin, which is essential for restoring organ-specific signal in liquid biopsy applications ^4,8,9^.

Fully harnessing the depth and breadth of EV-and other EP-derived multi-omic data in circulation could vastly expand the diagnostic and prognostic capabilities of liquid biopsy ^10–13^. This includes improving early disease detection, patient stratification, therapeutic monitoring, and addressing many other diagnostic gaps, especially in conditions where cfDNA-based approaches perform poorly. For instance, this is the case in cancers characterized by low cellular turnover and minimal ctDNA shedding, such as hepatocellular carcinoma (HCC), non-small cell lung cancer (NSCLC) with adenocarcinoma, brain tumors, and cancers of the thyroid, kidney, esophagus, head and neck, breast, and prostate, especially at early disease stages. ^14–16^.

Despite this promise, the clinical translation of EV biomarkers has been limited by the multicellular heterogeneity of plasma-derived EVs. ^17,18^.

Standard isolation methods such as ultracentrifugation, polymer precipitation and size-exclusion chromatography (SEC) recover mixtures of EVs, other EPs including lipoproteins and soluble proteins without distinguishing cellular origin. Even with the increasingly popular SEC method ^19–21^, information-rich tissue-derived EVs, which may reach up to 1 × 10⁵ EVs/mL of blood ^22,23^ are obscured by more than 1 × 10^9^ EVs/mL of blood-cell-derived EVs^20,24.^

This overwhelming background leads to false discoveries and spurious biomarker associations^25^ particularly in high-dimensional omics studies ^26^ and has hindered clinical translation. Targeted immunocapture of tissue-specific EVs using surface proteins indicative of cellular origin^27^ can reduce this noise and reveal clinically relevant pathways that remain masked in bulk EV analysis ^26^. Immunocapture is typically performed with antibody-coated magnetic beads (MBs)^28^.

However, most immunocapture studies have struggled to confirm the provenance of captured EVs^29–33^. Three critical challenges remain. First, ubiquitous EV markers such as the cytosolic Syntenin-1, Alix, TSG101 are poorly suited for selective EV capture and the commonly used tetraspanins CD9, CD63 and CD81 are not tissue-specific ^7,34–36^. Second, detection platforms such as nanoflow cytometry and ELISA often lack the sensitivity required to verify tissue enrichment on a targeted EV sub-type ^37,38^. Third, biomarkers that once seemed specific to a tissue-type are being debated, as highlighted with recent findings challenging the use of L1CAM as a marker to capture neuronal-EVs through plasma^39,40^.

Recent studies reporting enrichment of brain-, cardiac-, or adipose-tissue-derived EVs using RNA sequencing (RNA-seq) and/or proteomics show progress but still lack orthogonal validation such as surface protein comparison across tissues, analysis of cargo specificity and/or matched bulk EV analysis ^41,42,43,44,45,46^. These gaps underscore the need for workflows that confirm marker expression, antibody specificity and biological relevance of captured EV cargo.

In this study we introduce NEXPLOR (Novel EXtracellular vesicle PopuLation and Omics Revealer), a magnetic bead-based platform designed to selectively enrich tissue-derived EVs. NEXPLOR is applied here to isolate hepatocyte-derived EVs from bulk EVs derived from human plasma. We use three validation steps: (1) a bioinformatic pipeline that integrates transcriptomic, proteomic and subcellular localization data to select liver-specific capture markers; (2) rigorous antibody validation on EVs obtained from multiple snap-frozen tissues, quantified with O-NEXOS, an ultrasensitive EV detection platform; and (3) comprehensive downstream profiling of h-EV (hepatocyte-EVs) cargo.

To evaluate the performance of our platform, we first assessed its analytical performance against a similar commercial method for a known EV marker, CD9. After evaluation, we obtained a biomarker signature for h-EV capture and assembled a clinical cohort comprising two distinct patient subsets for h-EV isolation from bulk EVs in plasma samples. This cohort reflects a major diagnostic challenge for current standard-of-care approaches ^47,48^, including DNA-based liquid biopsy tests ^49,50,51^, in the early detection of HCC among patients with cirrhosis. The first subset is dedicated to h-EV proteomics analysis using timsTOF SCP, an LC-MS/MS system with single cell proteomics resolution. H-EVs showed strong enrichment of hepatocyte markers, depletion of common plasma contaminants, and the emergence of disease-related proteomic pathways not detected in bulk isolates. The second subset was analyzed for miRNA profiles. H-EVs revealed DE miRNAs known to be involved in HCC progression, many of which were undifferentiated in matched bulk EV samples. This addresses a longstanding limitation of miRNA-based liquid biopsies, namely the lack of tissue specificity.

Together, our findings establish NEXPLOR as a robust and scalable platform for isolating tissue-specific EVs from blood. This work sets a precedent for organ-specific EV profiling in plasma and expands potential to address longstanding limitations of bulk EV analysis. By restoring tissue specificity, NEXPLOR opens new opportunities for early disease detection, pharmacodynamic monitoring, and precision medicine applications.

## RESULTS

### 1. NEXPLOR efficiently isolates CD9^+^ EV subtypes without altering CD9 protein affinity

We compared the EV subtype immunocapture and isolation of NEXPLOR with an existing commercial kit, Exo-Flow^TM^. Similar to NEXPLOR, Exo-Flow allows for EV elution after capture, enabling direct performance comparison and validation of NEXPLOR against a reference method. To compare both methods, EPs isolated from MCF-7 and BT-474 cultures, which are known to express high levels of CD9 protein ^52^, were captured using Anti-CD9 antibody-coated NEXPLOR and Exo-Flow beads for CD9^+^EV isolation. The initial sample (starting sample), the supernatant from the isolation and isolated particles were characterized by Nanoparticle Tracking Analysis (NTA), for total EP concentration and O-NEXOS ^53^ for CD9^+^ EV detection across the three fractions (**Figure 1A**). As expected, the combined particle concentration of supernatant and isolated particle fractions is less than the concentration of the starting samples, as particles are expected to be lost during washing steps or due to incomplete particle elution from the beads. The extent of the loss could not be quantified. Exo-Flow yielded higher particle concentrations than NEXPLOR in both the supernatant and isolated fractions (**Figure 1B**). Illustrations in **Figure 1C** and **Figure 1D**, respectively, depict NEXPLOR for EV capture/isolation and O-NEXOS for detection in a sandwiched assay using biomarker reporters. Despite yielding lower particle concentrations, NEXPLOR effectively enriched or depleted CD9^+^ EVs, respectively, on the isolated particle and supernatant fractions but the isolated fraction of Exo-Flow did not contain CD9^+^ EVs, which remained in the supernatant (**Figure 1E**). To further assess the yield and specificity of CD9⁺ EV capture using NEXPLOR, we performed two rounds of NEXPLOR isolation (**Figure 1F**). NTA measurements showed that, on the second isolation round, NEXPLOR yielded a particle recovery of 97 % (**Figure 1G**) and the specific signal for CD9^+^ EVs, was enhanced over that of the starting sample, strongly suggesting a successful EV sub-type enrichment (**Figure 1H**). EV nominal size was not affected by successive isolations (**Figure S1**).

**Figure 1.**
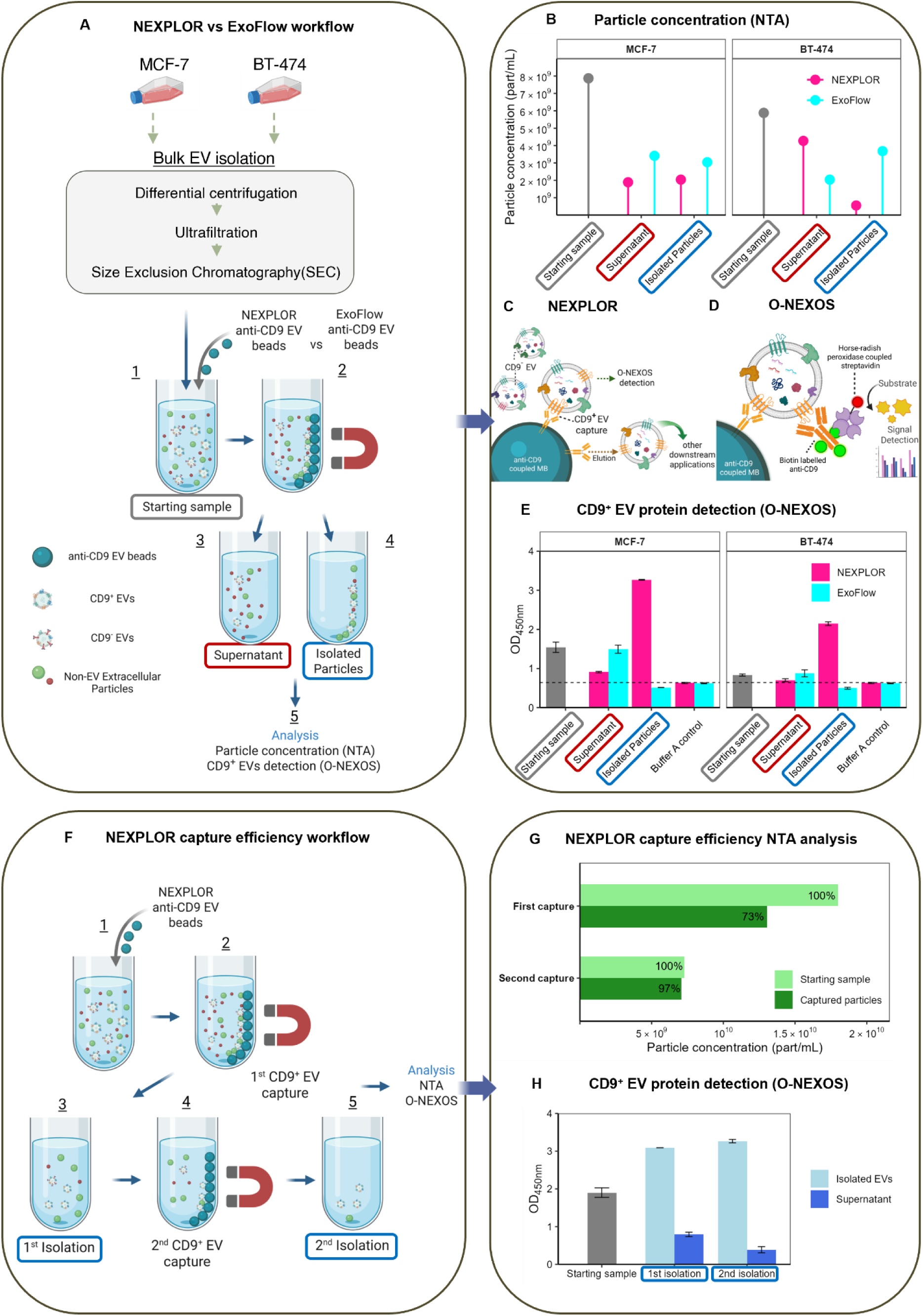
NEXPLOR effectively isolates a sub-population of CD9^+^EVs. **(A)** Workflow to compare the CD9^+^ EV isolation performance of the two immunocapture strategies. Cell culture supernatants from breast cancer cell lines (MCF-7 and BT-474) were cleared of cells and debris by centrifugation and then concentrated using ultrafiltration. The sample was further processed with SEC to isolate extracellular particles (1). Anti-CD9 NEXPLOR beads or anti-CD9 ExoFlow beads were used to capture CD9^+^ EVs (2) and then eluted (4). The supernatants with non-captured particles (3) and eluted CD9^+^ EVs fractions (4) were analyzed by nanoparticle tracking analysis for particle detection (NTA) and immunoassay for protein detection (CD9^+^ O-NEXOS). **(B)** Particle concentration of the Starting samples, Supernatant, and Isolated particles. Schematic of **(C)** NEXPLOR capture of EVs and **(D)** O-NEXOS detection of EVs via surface protein detection. **(E)** CD9^+^ O-NEXOS protein detection of the Starting samples, Supernatant, and Isolated particles after normalizing all the samples to 1x108 part/mL. The dotted line represents the Buffer A control mean OD450nm. **(F)** Workflow to verify the capture efficiency of the NEXPLOR anti-CD9 EV beads using the MCF-7 cell line only. After SEC (1, Starting sample) followed by CD9^+^ EV capture (2), the eluted fraction (3) was subjected to a second capture (4) with NEXPLOR anti-CD9 beads and eluted again (5). All supernatants with non-captured particles (not shown) and eluted CD9^+^ EVs fractions (3) (5) were analyzed by NTA for particle concentration and CD9^+^ O-NEXOS for EV protein detection. **(G)** NEXPLOR CD9^+^EV capture efficiency calculated by taking the sample after SEC isolation (Starting sample) minus the particles found in the supernatant after CD9^+^ EV capture (Captured particles). **(H)** CD9^+^ O-NEXOS EV protein detection of the Starting sample, Supernatant, and Isolated particles.

### 2. Bioinformatic Workflow Guides Identification of Hepatocyte-Specific EV Capture Markers

Having validated NEXPLOR for the capture and isolation of CD9^+^ EVs, we aimed to adapt NEXPLOR to isolate hepatocyte-derived EVs. The isolation of h-EVs may be a strategy to facilitate cargo enrichment for clinical applications in liver disease liquid biopsies. ASGR1 and ASGR2 are commonly recognized as hepatocyte-specific markers and, ASGR1 has been previously targeted for the isolation of hepatocyte-derived EVs ^54^. However, ASGR1 and ASGR2 are also expressed at lower levels in peripheral blood monocytes and ASGR1 is often downregulated in HCC, limiting their universal use for global h-EV isolation^55,56^.

This motivated our search for finding additional biomarker candidates to capture and isolate h-EVs. For that, we developed a bioinformatics workflow to guide our selection. First, by querying the Human Protein Atlas (HPA), we identified 936 genes with elevated mRNA expression in liver compared to non-liver tissues. From this initial list, we filtered out 552 genes with known expression at the protein level, defined as “high” and “medium” expression in non-liver tissues, resulting in 384 protein target candidates. Secondly, we identified genes with “high” and “medium” expression at the protein level in hepatocytes and filtered out those also presenting “high” and “medium” protein expression in non-liver tissues, resulting in a list of 70 proteins. From the combined total of 413 proteins, we narrowed our candidates to 81 that are primarily localized to the plasma membrane. We hypothesize that some of these may decorate the surface of hepatocyte-derived EVs (**Figure 2A; full list in Table S1**).

**Figure 2.**
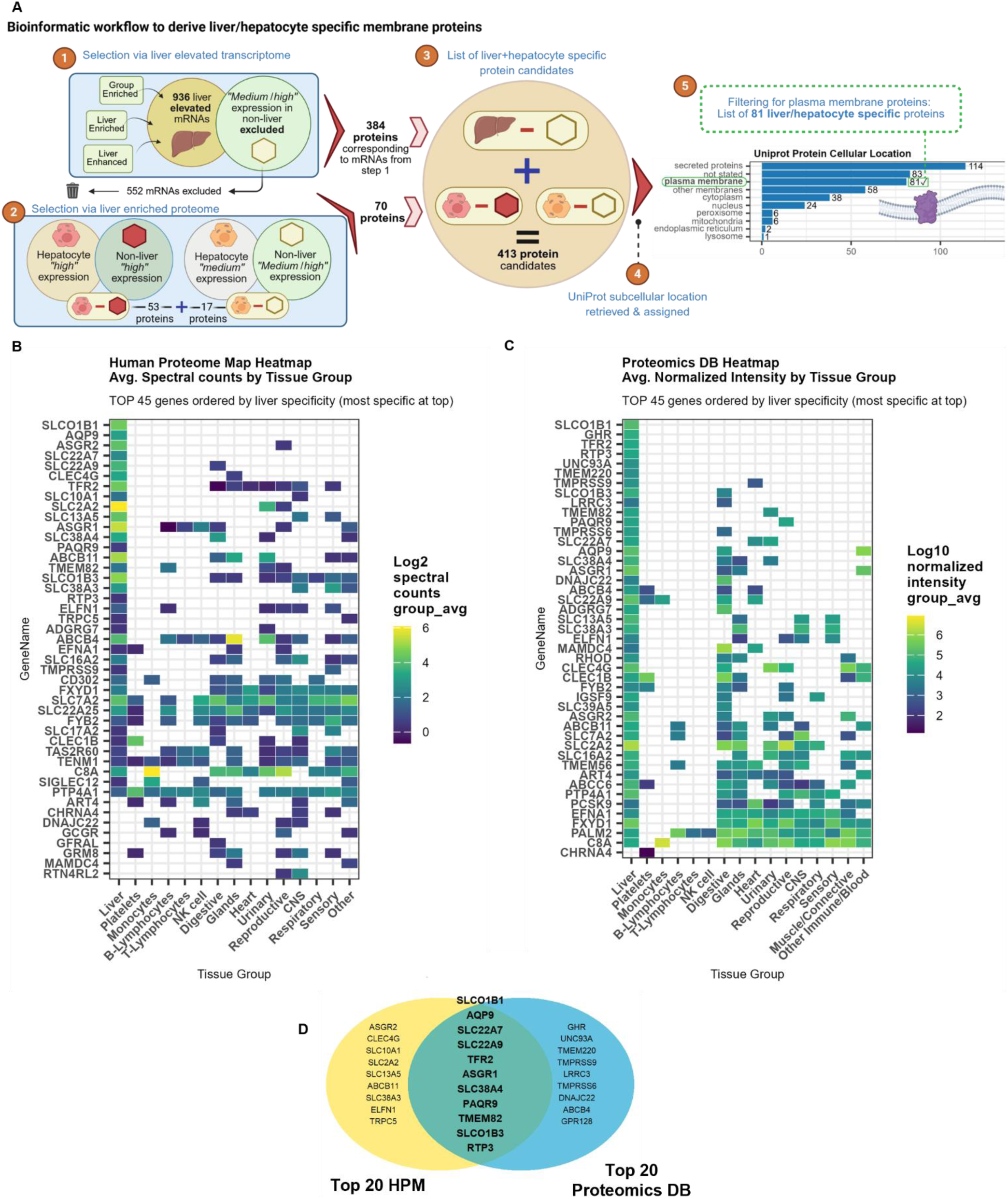
Data mining of publicly available datasets to identify and verify liver- and hepatocyte-specific plasma membrane proteins. **(A)** Summary of bioinformatic pipeline applied to mRNA and protein datasets within the Human Protein Atlas (HPA) to generate a combined hepatocyte and liver-specific protein list. Proteins in this list were excluded if they showed high or medium expression in non-liver tissue. The resulting 413 protein candidates were then mapped to subcellular locations as annotated on the UniProt database. 81 proteins were localized to the plasma membrane for further analysis. **(B)** Heatmap shows log₂-transformed spectral counts (mass spectrometry) for the top 45 markers, ranked by liver specificity, with most specific candidates on top, out of the 81 candidate proteins and their relative expression in various organs. Data was collected from the Human Proteome Map (HPM). Data was not available for 29 protein candidates. **(C)** Heatmap shows log10-normalized intensity values (mass spectrometry) for the top 45 markers, ranked by liver specificity with most specific candidates on top, out of the 81 candidate proteins and their relative expression in various organs. Data collected from the Proteomics DB database. Data was not available for 27 protein candidates. **(D)** Venn diagram displaying the overlap of the top 20 proteins ranked by liver specificity in both HPM and Proteomics DB rankings. Yellow: 9 candidate proteins that are top20 exclusively for the HPM ranking, Green: 11 Protein candidates that are on the top 20 for both rankings, Blue: 9 candidate proteins that are top 20 exclusively for the Proteomics DB ranking.

To evaluate their tissue specificity, we cross-referenced these 81 proteins against mass-spectrometry profiles from multiple tissues in the Human Proteome Map (HPM) (**Figure 2B**) and ProteomicsDB (**Figure 2C**). In each database, we ranked the proteins by a liver specificity ratio, calculated as liver intensity divided by intensity in non-liver tissues, and the top 45 candidates are shown in **Figures 2B** and **2C**, respectively. Markers identified as highly liver-specific in both databases were prioritized for validation. However, we also considered candidates that ranked highly in only one database elevating our final candidate selection to 29 proteins (**Figure 2D**). For instance, ASGR2 was the third most liver-specific protein in HPM but did not rank among the top 20 in ProteomicsDB. Tissue-specific data was unavailable either in one or both databases for 27 of the 81 proteins in the initial list.

### 3. NEXPLOR preferentially enriches hepatocyte-derived EVs from human tissues

Of the 29 shortlisted protein targets through the workflow in **Figure 2**, 10 lacked antibody compatibility with NEXPLOR and O-NEXOS due to insufficient analogous ELISA or IP validation. To expand options, four additional markers (ABCC6, PTP4A1, SLC17A4 and TMEM56) with strong antibody compatibility with NEXPLOR and O-NEXOS were included. Antibodies were coupled to NEXPLOR MBs and validated via CD63-based O-NEXOS, as it was the only tetraspanin we detected in HepG2 EVs (**Figure S2**) and, according to HPA, it is preferentially enriched in the liver compared to e.g. CD9 or CD81 (**Figure S3**), other commonly used tetraspanins in EV studies^57^.

In a first validation step aimed to detect unspecific cross-reactions, sixteen candidates cross-reacted with antibody anti-CD63 and were excluded from further analysis, narrowing the list to 13 markers. Detected cross-reaction levels were above our in-house developed cut-off of 0.4 OD450nm (Optical Density) in buffer-only controls (**Table S2**). Selected 13 antibody-coupled MB types were then incubated with the EV preparations (including EP types) derived from HepG2 cell culture and HCC tissue (see Materials & Methods), for CD63-based O-NEXOS detection. Based on assay performance, our selection was further narrowed to 10 protein targets, namely ASGR1, ASGR2, TFR2, SLCO1B1, FXYD1, SLC2A2, SLC22A9, TMEM56, SLC38A3, and UNC93A.These were selected based on strong signal-to-noise (S/N) ratio and a minimum difference of 0.5 OD units at 450 nm (OD450nm) between signal and background (**Table 1**). To assess the NEXPLOR specificity for detecting hepatocyte-specific biomarkers, we evaluated the 10 selected antibodies using EV preparations derived from both liver tissues (healthy and HCC) and non-liver tissues (kidney and lung) (**Figure 3A** and **Table S3**). From this pre-screening, we selected the antibodies that showed an S:N ratio greater than 2 exclusively in liver-derived EVs. This analysis led to the final selection of four antibodies targeting the proteins ASGR1, ASGR2, TFR2, and SLCO1B1, collectively referred as the h-EV TOP4 capture panel (highlighted in **Figure 3A**). The performance of each TOP4 antibody individually and in combination as a single capture antibody cocktail (**Figure 3B**), was evaluated across an expanded panel of tissue-derived EVs, including both liver tissues (healthy, HCC, cirrhosis) and non-liver tissues (kidney, lung, breast, adipose, muscle). All samples were normalized to a final concentration of 1x10^10^ particles/mL. Notably, double-positive ASGR1^+^CD63^+^, ASGR2^+^CD63^+^ and TFR2^+^CD63^+^ EVs were consistently detected at high levels across all liver tissues, with signal intensities approaching assay saturation, for ASGR1^+^CD63^+^. Signal was considerably reduced for non-liver tissues (**Figure 3C**).

**Figure 3.**
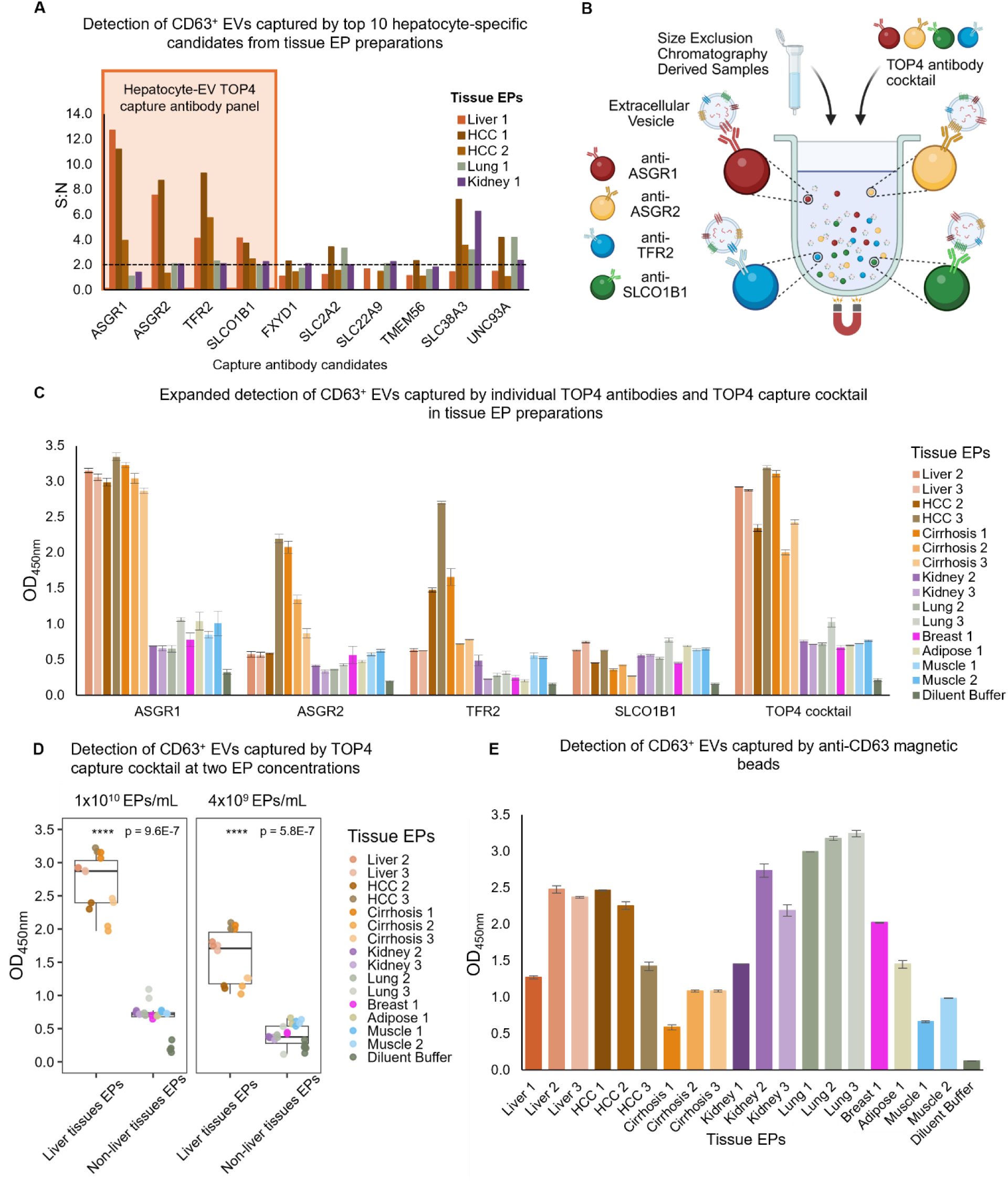
Selection and characterization of the NEXPLOR hepatocyte-EV TOP4 capture antibody panel using O-NEXOS. **(A)** Signal-to-noise (S:N) ratios following CD63^+^ EV detection for each of the top ten final candidate capture antibodies were calculated by dividing the signal detected in tissue extracellular particles (Tissue EPs) by the signal in diluent buffer controls. Tissue EPs from healthy liver (Liver), diseased liver (HCC) and non-liver tissues (lung, kidney) were examined to evaluate hepatocyte specificity. Antibodies with a S:N ratio greater than 2 (dotted line) exclusively in liver-derived EVs were selected for the hepatocyte-EV (h-EV) TOP4 capture panel, targeting collectively the proteins ASGR1, ASGR2, TFR2 and SLCO1B1, as highlighted in the pink-shaded region. **(B)** Schematic representation of the h-EV TOP4 capture antibodies anti-ASGR1, anti-ASGR2, anti-TFR2 or anti-SLCO1B1 coupled to magnetic beads (MBs) and combined to make the TOP4 capture cocktail for multiplexed EV capture. **(C)** Optical density at 450 nm (OD_450nm_) from CD63^+^ EV detection following capture with each individual TOP4 antibody and with the combined TOP4 cocktail, in extracellular particles (EPs) derived from liver and non-liver tissues. All samples were normalized to 1x10^10^ particles/mL. Data are presented as mean ± standard error of the mean (SEM), n = 2. **(D)** CD63^+^ EVs detection at OD_450nm_ using the TOP4 capture cocktail, in liver and non-liver tissues EPs at two concentrations: 1x10^10^ and 4x10^9^ EPs/mL. Statistically significant differences were observed between liver and non-liver tissue EPs at both concentrations (Kruskal-Wallis test, 1x10^10^ EPs/mL: p = 9.6E-7; 4x10^9^ EPs/mL: 5.8E-7). Data represents n=2 per tissue EP samples, and Diluent Buffer 1x10^10^ EPs/mL: n=4, 4x10^9^ EPs/mL: n=6. **(E)** CD63^+^ EV detection at OD_450nm_ in liver and non-liver derived EPs normalized to 5x10^8^ particles/mL following capture with anti-CD63 magnetic beads (MBs) as a control. No statistically significant differences were observed between liver and non-liver tissue EPs (Kruskal-Wallis test, p = 0.3267). Data are presented as mean ± SEM, n = 2.

**Table 1.** O-NEXOS screening results for NEXPLOR antibody candidates*

|  | HepG2 EVs |  | HCC EVs |  |
| --- | --- | --- | --- | --- |
| Protein | S-N | S:N | S-N | S:N |
| ABCC6 | 0.09 | 1.59 | 0.16 | 2.13 |
| **ASGR1** | **3.11** | **29.89** | **0.22** | **3.05** |
| **ASGR2** | **1.30** | **11.29** | **2.99** | **17.88** |
| ELFN1 | 0.36 | 2.32 | 0.22 | 1.87 |
| **FXYD1** | **2.09** | **6.05** | **1.21** | **2.94** |
| SLC17A4 | 0.43 | 0.56 | 0.23 | 2.15 |
| **SLC22A9** | **0.33** | **2.04** | **0.80** | **2.72** |
| **SLC2A2** | **0.19** | **2.43** | **1.09** | **3.91** |
| **SLC38A3** | **2.70** | **7.64** | **1.09** | **2.29** |
| **SLCO1B1** | **1.04** | **6.91** | **0.97** | **4.74** |
| **TFR2** | **0.18** | **2.56** | **1.58** | **6.98** |
| **TMEM56** | **1.63** | **1.63** | **0.98** | **4.85** |
| **UNC93A** | **1.59** | **1.59** | **1.01** | **6.19** |

\*Proteins, in bold, were selected for further validation and showed at least a two-fold increase in O-NEXOS signal relative to background (signal-to-noise ratio) and a minimum difference of 0.5 Optical Density (OD) units at 450 nm ( $OD_{450nm}$ ) between signal and background

SLCO1B1 demonstrated less specificity than expected but based on the workflow in **Figure 2** and initial experimental data, we opted to include it into our h-EV TOP4 capture antibody cocktail. The antibody cocktail was tested at a normalized particle concentrations of 1x10^10^ and 4x10^9^ particles/mL for all tissue-derived EVs. At both tested concentrations (1x10^10^ and 4x10^9^ particles/mL), the TOP4 capture antibody cocktail produced significantly higher CD63^+^ signals in EVs derived from healthy liver, cirrhotic and HCC tissue samples compared to those derived from other tissue sources (**Figure 3C** and **Figure 3D**). These differences were statistically significant, as determined by Kruskal–Wallis tests ^58^ (p = 9.6×10⁻⁷ and p = 5.8×10⁻⁷ at 1×10¹⁰ and 4×10⁹ particles/mL, respectively; *n* = 2 per tissue EP sample), reinforcing the hepatocyte specificity of the selected TOP4 capture antibody panel. To further validate the specificity of our detection approach, EPs derived from the various tissue sources, were also analyzed for CD63^+^ EVs capture and detection with O-NEXOS (**Figure 3E**). No statistically significant differences were observed between liver and non-liver EPs (Kruskal–Wallis test, p = 0.3267; normalized to 5×10⁸ particles/mL, n = 2), indicating that CD63 is not preferentially expressed in hepatocyte-derived EVs. This rules out CD63 as a confounding factor and confirms that the enhanced signals observed with the TOP4 antibodies are due to the specific or enriched expression of target biomarkers on hepatocyte-originating EVs. These findings confirm selectivity of the TOP4 capture antibody panel for detecting hepatocyte-derived EVs using the NEXPLOR platform.

### 4. NEXPLOR Enables Deep and Reproducible Proteomic Profiling of Hepatocyte-Derived EVs compared to Bulk EVs in Clinical Plasma Samples

Since specificity to circulating EVs from a given tissue is virtually impossible to ascertain from bulk EV preparations, we developed the validation using EV preparations locally derived from several snap-frozen tissues, as demonstrated in the previous section, but, with the objective to translate the methodology to bulk EV preparations from human plasma samples. With this goal, we first validated the h-EV capture cocktail for plasma unspecific signal by evaluating the performance of the h-EV cocktail against an control mixture of MBs made with isotype antibodies. The S:N ratio was always higher than 1.0 demonstrating that the h-EV capture cocktail preferentially captures EVs that can be detected by via CD63 signal (**Figure S4A**), as previously shown (**Figure 3**). Demographic details of samples used are available in **Table S3**.

To determine the advantage of the h-EV capture method over bulk EV isolations, we assembled a small clinical cohort comprised of 13 HCC with underlying cirrhosis (HCC) patient samples, 10 cirrhosis without HCC (Cirrhosis) samples and 6 healthy donors (Healthy). Clinical metadata is available in **Table S4**. The 29 samples were processed for bulk EV isolation using SEC. The final elution volume for each sample was split into two aliquots: one was retained for analysis as a bulk EV isolate, and the other was processed using the NEXPLOR method to isolate hepatocyte-derived EVs. The resulting samples were processed for LC-MS/MS, in a paired design experiment (**Figure 4A**).

**Figure 4.**
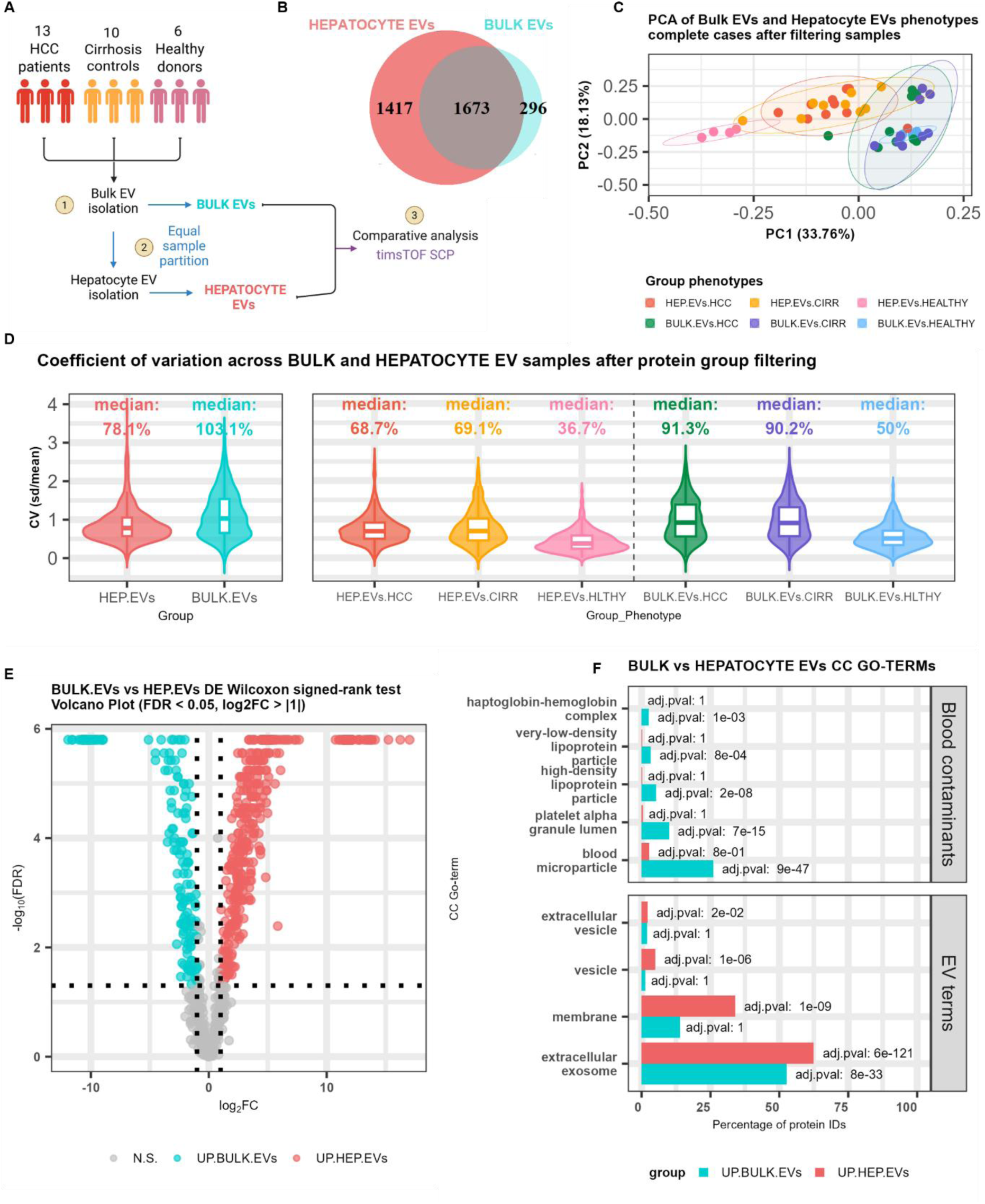
**Exploratory proteomic and differential abundance (DE) analysis across hepatocyte and bulk EV preparations**. **(A)** Schematic workflow for EV isolation and comparative proteomic analysis using timsTOF SCP. Plasma samples were collected from HCC patients (n = 13), cirrhosis controls (n = 10), and healthy donors (n = 6). EVs were first isolated as bulk EVs, then samples were equally partitioned with hepatocyte EV (h-EV) isolation being applied to one portion to be analyzed by LC-MS/MS with timsTOF SCP. **(B)** Venn diagram displaying the overlap of protein group (PG) identifications from MS measurements of hepatocyte and bulk EV samples. Red: 1417 PGs uniquely identified in hepatocyte EV samples; blue: 296 PGs uniquely identified in bulk EV samples; grey: 1673 PGs identified in both. **(C)** Principal component analysis (PCA) using complete cases before missing-value imputation, with samples color-coded by group and phenotype: Hepatocyte EVs (h-EVs) – red (HEP.EVs.HCC), orange (HEP.EVs.CIRR), pink (HEP.EVs.HEALTHY); Bulk EVs – green (BULK.EVs.HCC), purple (BULK.EVs.CIRR), light blue (BULK.EVs.HEALTHY). Confidence ellipses (95%) were computed using a multivariate t-distribution, with Euclidean distance. **(D)** Violin plots and overlaid boxplots representing the coefficient of variation (CV) of PGs intensity values across samples, after PG filtering. The left panel compares hepatocyte (red) and bulk (blue) EV samples. The right panel further stratifies by group and phenotype: same color scheme as in (C). CV is shown on the Y-axis; median CV values are annotated as percentages above each group. **(E)** Volcano plot of DE analysis results of bulk vs hepatocyte EV samples. PGs significantly enriched in bulk EV samples are colored in light blue, and PGs significantly enriched in hepatocyte EV samples are colored in red, considering an adjusted p-value threshold of 0.05 and a log2 fold-change (log₂FC) of 1. **(F)** Comparative functional enrichment analysis of Gene Ontology (GO) cellular component (CC) terms related to blood contaminants in PGs differentially abundant (DE) between bulk and hepatocyte EV samples. The x-axis indicates the percentage of upregulated PGs annotated to each CC term for each sample type, and Benjamini–Hochberg adjusted p-values are overlaid as text labels.

The experiment was initially planned for 30 plasma samples, but from the 30 pairs of hepatocyte EV and bulk EV samples, 1 pair of cirrhosis samples failed during data acquisition due to technical issues. An additional six samples (2 Healthy, 1 cirrhosis and 3 HCC) were excluded based on quality control criteria, specifically having total protein group (PG) counts below the 10th percentile across all samples. Their respective matched samples were also removed to maintain the paired design. After sample filtering, 23 matched pairs (23 hepatocyte EV and 23 bulk EV preparations) were retained for downstream analysis.

Across these, a total of 3386 PGs were quantified, of which 1417 PGs were exclusive to hepatocyte EV preparations (**Figure 4B**). Following the removal of low abundant PGs, 859 PGs remained, of which 47 were unique to hepatocyte EV samples and 35 to bulk EV samples (**Figure S4B**). Clustering by Principal Component Analysis (PCA) shows that h-EV samples from healthy donors were tightly clustered, with their respective multivariate t-distribution 95% confidence interval (CI) ellipse only briefly overlapping those of liver disease (HCC and cirrhosis) patients indicating that healthy donors’ samples remain discernible in multivariate space **(Figure 4C**, **Figure S4C**). We monitored the h-EV and Bulk EV phenotypes separation along the procedures of our data processing and filtering (**Figure S4D to S4G**), Technical reproducibility for h-EV isolations was higher than bulk EVs, both before and after the PG filtering step. Additionally, all groups displayed an increase in coefficient of variation(CV%) after PG filtering, but more predominant in bulk EVs (**Figure 4D** **and Figure S4H**). See Materials and Methods.

Differential expression analysis (DE) identified PGs differentially expressed, or “abundant”, in bulk and h-EVs, with 366 PGs significantly more abundant in h-EV samples, and 181 PGs in bulk EVs. Statistical significance was defined as an adjusted p-value < 0.05 and an absolute log₂ fold change > 1 (**Figure 4E** **and Table S5**).

Proteins enriched in hepatocyte-derived EVs showed stronger association with EV- and membrane-related GO terms than those in bulk EVs (e.g., 62.4% vs. 52.7% for EV-related GO:0070062). In contrast, bulk EVs were notably enriched for blood contaminants, including markers of blood microparticles, lipoproteins, and platelet granules, and signals largely absent in NEXPLOR isolates (**Figure 4F**).

### 5. NEXPLOR-Based h-EV Enrichment Reveals EV, Liver-Specific and Metabolic Protein Signatures with reduced Contaminant Background

We further validated our proteomics data using the MISEV2023 EV characterization framework (**Table S5**) ^57^. We noted that several PGs belonging to the categories “1-Transmembrane (or GPI-anchored) proteins associated with plasma membrane and/or endosomes” and “2-Cytosolic proteins in EVs” were more abundant for h-EVs compared to bulk EV samples. EV proteins such as LAMP1, LAMP2, CD82 and several tubulin proteins were significantly more abundant in h-EV samples while others such as PDCD6IP, CD81 and TFR2 showed positive log2 fold-change (log₂FC) did not meet the defined significance thresholds. In contrast several lipoproteins (category 3a NonEV lipoproteins) and PGs associated with the category “5 – Secreted proteins recovered with EVs” were found to be more significantly abundant in bulk EV samples (**Figure 5A** and **Table S5**). Furthermore, when evaluating the level of expression by heatmap prior to filtering PGs and IgGs it is noted that several IgGs are detected more often in bulk than h-EV isolations (**Figure S5**).

**Figure 5.**
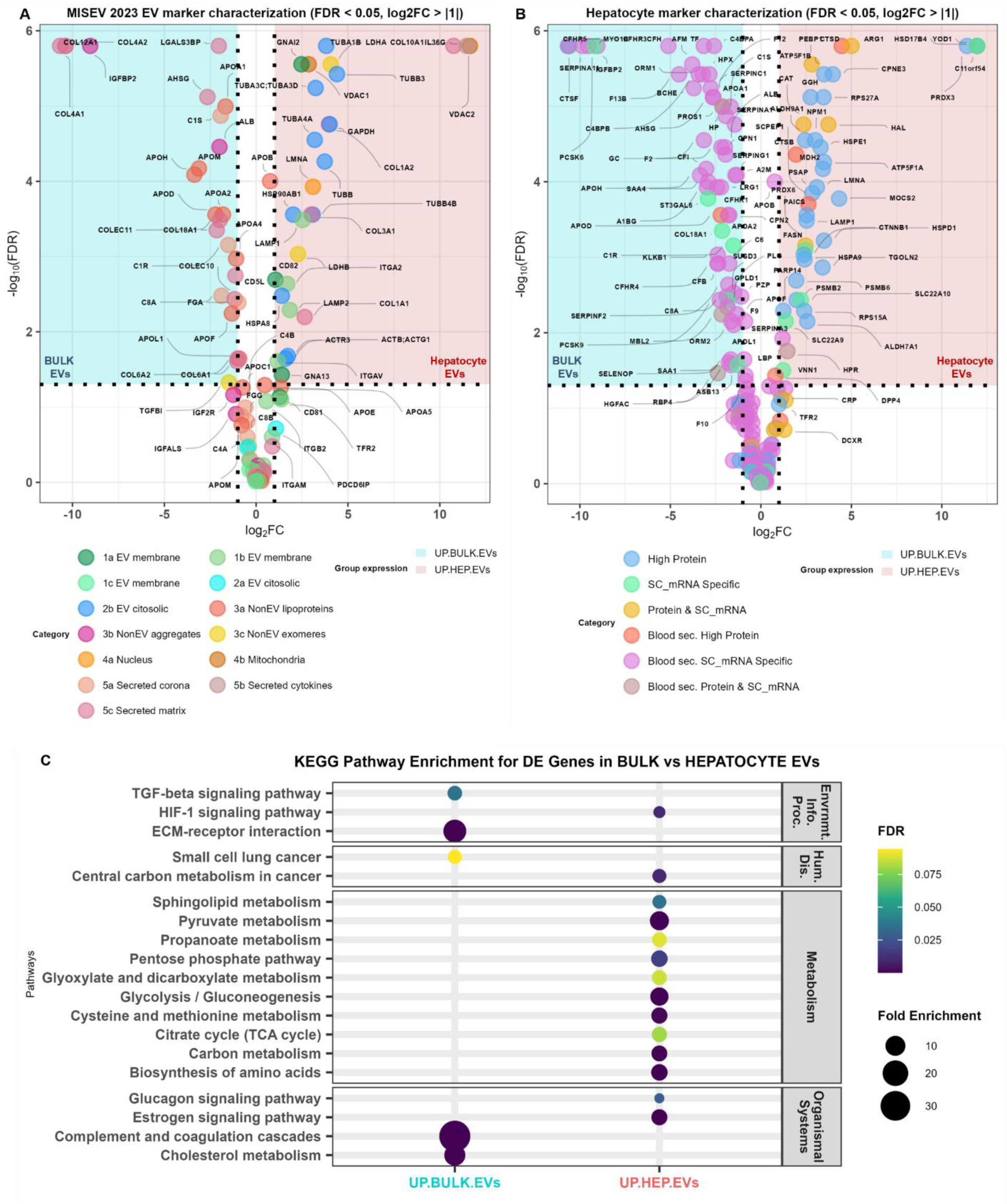
Paired differential abundance (DE) of EVs markers, EV co-contaminants and hepatocyte markers and KEGG pathways associated with the enriched proteome in bulk and hepatocyte EV samples. (A) Volcano plot of DE results after bulk EV or NEXPLOR isolation, with PGs classified according to the MISEV2023 content-based EV categories: 1 a-c (green hues): Transmembrane (or GPI-anchored) proteins associated with plasma membrane and/or endosomes; 2 a-b (blue hues): Cytosolic proteins in EVs; 3 a-c (red, orchid, yellow): Major components of non-EV co-isolated structures (NVEPs); 4 a-b (orange, brown): Transmembrane, lipid-bound and soluble proteins associated with intracellular compartments other than plasma membrane/ endosomes; 5 a-c (pink hues): Secreted proteins recovered with EVs; PGs enriched in NEXPLOR are shown in the right, red-shaded area (positive log₂FC); PGs enriched in bulk EV are shown in the left, blue-shaded area (negative log₂FC). **(B)** Volcano plot of differentially expressed (DE) results after bulk EV or NEXPLOR isolation, focusing on PGs with high protein expression and/or specific mRNA expression in hepatocytes, based on annotations from the Human Protein Atlas (HPA). Dots (PGs) are color-coded and annotated as follows: High Protein (blue): High protein expression in hepatocytes; SC_mRNA Specific (light green): Hepatocyte-specific expression based on single-cell RNA-seq; Protein & SC_mRNA (gold yellow): Both high protein and SC mRNA expression in Hepatocytes; Blood sec. High Protein (red): High hepatocyte protein expression, also secreted into blood; Blood sec. SC_mRNA Specific (purple): Hepatocyte-specific by SC mRNA and secreted into blood; Blood sec. Protein & SC_mRNA (brown): Secreted into blood and meets both expression criteria PGs enriched in NEXPLOR (UP_NEXPLOR) are shown in the right, red-shaded area (positive log₂FC); PGs enriched in SEC (UP_SEC) are shown in the left, blue-shaded area (negative log₂FC). **(C)** Dot plot showing significantly enriched KEGG pathways (Benjamini-Hochberg, BH, adjusted p-value < 0.1,) for protein groups (PGs) enriched in bulk EVs (UP.BULK.EVs) and hepatocyte EVs (h-EVs) (UP.HEP.EVs) samples. Dot color indicates false discovery rate (FDR), and size reflects fold enrichment.

Beyond EV-associated proteins, markers for non-EV co-isolated structures, such as exomeres and supermeres, specifically LDHA and LDHB, were more abundant in h-EV samples. Interestingly, mitochondrial proteins VDAC1 and VDAC2 were also detected almost exclusively in the h-EV samples. (**Figure 5A**, **Figure S5**).

Heterogeneous nuclear ribonucleoprotein A2B1 (HNRNPA2B1), an RNA binding protein that plays a key role for the loading of specific miRNA into EVs was enriched in h-EV samples with a log₂FC of 2.51. Other members of the hnRNP family (HNRNPK and HNRNPH1) and ANXA2, involved in the sorting of mRNA or miRNA into EVs^59,60,61^ were also enriched.

Finally, a heatmap evaluation of EV sub-population markers, as described by Kowal et al.^62^, showed that these markers were more prominent in h-EV samples compared to bulk EVs samples. This was particularly evident for markers associated with CD63-positive EVs (**Figure S6**).

We next examined HPA annotated PGs as having high hepatocyte protein expression or hepatocyte-specific mRNA expression and compared their abundances between bulk and h-EV samples. Both positive and negative log₂ fold-changes were observed, indicating that each method enriches distinct subsets of hepatocyte-related proteins. However, most PGs enriched in bulk EV samples (negative log₂FC) belonged to the blood secretome category, whereas most PGs enriched in hepatocyte EV samples (positive log₂FC) did not fall into this classification (**Figure 5B**). Additionally, most of the PGs annotated as having both specific single-cell mRNA expression and high protein expression in hepatocytes, but not classified as actively secreted into blood circulation, were enriched by NEXPLOR. Interestingly, samples clustered perfectly into bulk and h-EV sample groups when clustering based on the level of expression of these markers, before the PG filtering step (**Figures S7**).

KEGG pathway analysis performed on the 366 protein groups significantly enriched in hepatocyte EV samples (adjusted p-value < 0.05, |log₂FC| > 1) revealed a strong overrepresentation of liver-relevant metabolic pathways, including pyruvate metabolism, carbon metabolism, biosynthesis of amino acids, and glycolysis/gluconeogenesis ^63,64^, as well as the HIF-1α signaling pathway, which is implicated in liver disease ^65^. In contrast, the 181 PGs more abundant in bulk EV samples were enriched for extracellular matrix–receptor interaction, cholesterol metabolism, and complement and coagulation cascades pathways, the latter showing a considerable fold-enrichment of 33.8 (**Figure 5C**). The full list and further details of the KEGG pathways significantly enriched for each subset can be found on **Table S6**.

### 6. NEXPLOR isolation of hepatocyte extracellular vesicles from liver disease patients enriches for markers associated with HCC and cirrhosis relevant pathways

After applying the same filtering criteria and analytical pipeline used for the full cohort of 23 bulk and h-EV sample pairs to the disease-stratified subgroups, 13 pairs from patients with HCC and 10 pairs from patients with cirrhosis-only (cirrhosis), distinct method-dependent proteomic profiles were observed. When comparing all four liver disease subgroups together (h-EVs HCC, bulk EVs HCC, h-EVs cirrhosis and bulk EVs cirrhosis), 35 PGs were unique to h-EVs HCC, 69 to h-EVs cirrhosis, 10 to bulk EVs HCC, and 33 to bulk EVs cirrhosis samples (**Figure 6A**). The 35 h-EVs HCC NEXPLOR-specific PGs formed a statistically significant protein-protein interaction (PPI) network (PPI enrichment *p* < 1.0E–16) (**Figure 6B**), with enrichment in:

- GOCC: *extracellular exosome* (false discovery rate, FDR = 1.90E–09), *vesicle* (FDR = 1.24E–07);
- GOMF: *RNA binding*, *nucleic acid binding*;
- Reactome: Formation of a pool of free 40S subunits (FDR = 2.28E-08) and L13a-mediated translational silencing of Ceruloplasmin expression (FDR = 2.34E-08);
- Tissues: *liver* and *digestive gland* (FDR ≤ 1.09E–16).

**Figure 6.**
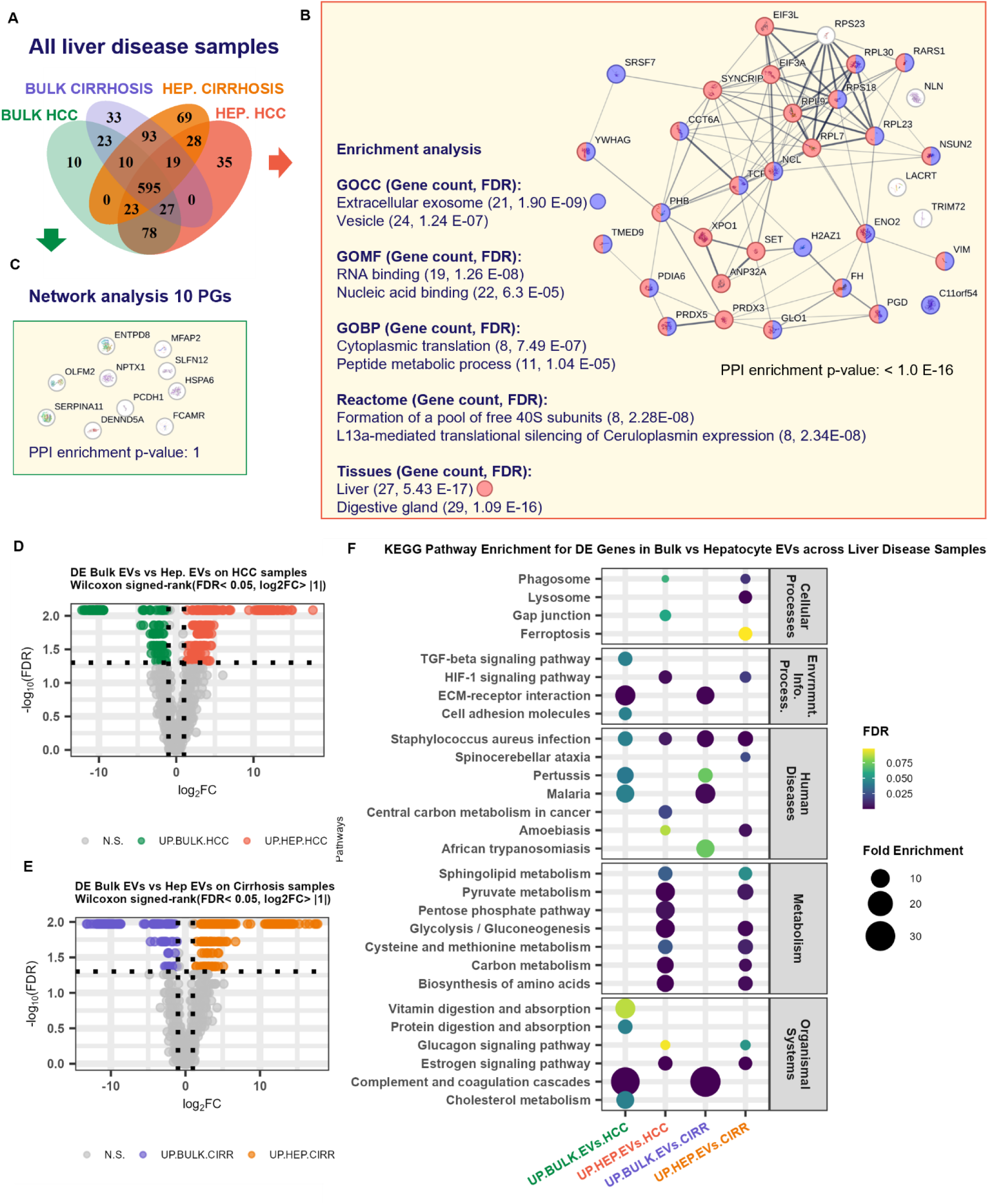
Hepatocyte and bulk EV comparative analysis in liver disease patient samples reveals that hepatocyte EVs enriches for liver disease related pathways. **(A)** Venn diagram showing PG overlap across all liver diseases (HCC and cirrhosis) in hepatocyte and bulk EV samples. **(B)** STRING protein-protein interaction (PPI) network analysis of the 35 PGs uniquely identified in hepatocyte EV HCC samples (red) compared to HCC and cirrhosis bulk EV (green and purple) and NEXPLOR cirrhosis samples (orange). Enrichment analysis was performed using Gene Ontology cellular component (GOCC), molecular function (GOMF), biological process (GOBP), Reactome, and Jensen Tissues databases. The two top enriched terms for each category are shown (right). PGs annotated with the GOCC term “extracellular exosome” are colored in purple, and those associated with the Jensen Tissues term “liver” are colored in red. **(C)** PPI network of the 10 PGs uniquely identified in bulk EV HCC samples compared to all NEXPLOR samples and bulk EV cirrhosis samples. No significant functional enrichment was detected. **(D-E)** Volcano plot of differential abundant (DE) analysis results of hepatocyte EV vs bulk EV samples segmented by phenotype: HCC with cirrhosis **(D)** and cirrhosis-only **(E)** patient samples. PGs significantly enriched for bulk EV samples are colored in green/purple, and PGs significantly enriched in hepatocyte EV samples are colored in red/orange, considering an adjusted p-value threshold of 0.05 and a log2 fold-change (log₂FC) of 1. **(F)** Dot plot showing significantly enriched KEGG pathways (Benjamini-Hochberg adjusted p-value < 0.1,) for PGs enriched in bulk and hepatocyte EV samples of patients with HCC and cirrhosis (UP.BULK.EVs.HCC and UP.HEP.EVs.HCC) and patients with cirrhosis-only (UP.BULK.EVs.CIRR and UP.HEP.EVsCIRR). Dot color indicates false discovery rate (FDR), and size reflects fold enrichment.

In contrast, the 10 PGs uniquely identified in HCC bulk EV samples formed no detectable PPI network and showed no statistically significant enrichment, suggesting a lack of functional or tissue coherence (**Figure 6C**).

We also compared the same bulk and h-EV samples for the same liver disease sub-group (**Figure S8**). For the HCC group, 82 PGs were uniquely identified in h-EVs compared to 43 in bulk EVs (**Figure S8A**). Functional enrichment analysis of the 82 PGs unique to h-EVs compared to bulk EVs in HCC samples revealed significant associations with extracellular vesicle biology, RNA metabolism, and liver tissue specificity (**Figure S8B**). In contrast, these terms were not significantly enriched for the bulk EVs uniquely identified PGs, underscoring the differences in EV populations captured by the two isolation methods (**Figure S8C**). Two proteins with well-established roles in miRNA sorting into EVs,HNRNPA2B1 and SYNCRIP,, the latter identified as a component of the hepatocyte exosomal miRNA sorting machinery, were exclusively identified in the NEXPLOR-HCC subset ^59,60,66^.

Similarly, 120 PGs were uniquely identified in h-EVs from cirrhosis samples versus 83 in bulk EVs (**Figure S8D**). For the cirrhosis-only group, “*extracellular exosome”* was enriched in both h-EVs and bulk EV subsets but with higher statistical significance and a larger gene count for NEXPLOR (51 PGs, FDR = 1.88E–16) versus SEC (26 PGs, FDR = 1.60E–04).

Importantly, only hepatocyte EV cirrhosis samples showed enrichment for the liver term (34 PGs, FDR = 2.37E–05), while bone marrow cell and plasma cell terms were again enriched in the SEC-specific subset, mirroring the pattern observed in HCC (**Figure S8E and S8F**). Interestingly, VDAC2 and VDAC3, two outer mitochondrial membrane proteins associated with liver disease and HCC^67,68^ were uniquely identified in hepatocyte EV samples from the cirrhosis-only group. The lists of unique identified PGs can be found on **Table S7**. Overall, these findings are consistent with the broader dataset, where NEXPLOR consistently captured a more diverse and liver-relevant subset of circulating proteins.

DE analysis was performed across liver disease phenotypes to capture additional difference into biological processes with the two EV isolation strategies. Volcano plots comparing bulk and h-EVs in HCC (**Figure 6D**) and cirrhosis (**Figure 6E**) samples revealed a substantial number of PGs with significantly altered abundance (FDR < 0.05, |log₂FC| > 1). KEGG pathway enrichment analysis on these DE PGs identified distinct pathways associated with each isolation method. In line with observations from the full cohort, several metabolic pathways and the HIF-1 signaling pathway were significantly enriched in h-EVs sample from both HCC and cirrhosis subsets compared to their bulk EVs counterparts.

Notably, the Ferroptosis pathway, implicated in chronic liver injury and fibrosis^69^, was significantly enriched in PGs upregulated in h-EVs from cirrhosis samples. In HCC, Central carbon metabolism in cancer, a pathway extensively characterized in liver tumor biology^70,71,72^, was specifically enriched for h-EVs from the HCC samples subset (**Figure 6F**). The full list of KEGG pathway enriched in this analysis can be found in **Table S8**. Together, these findings indicate that NEXPLOR preferentially captures disease-relevant biological pathways in both HCC and cirrhosis, offering improved resolution of pathophysiological processes compared to bulk EV isolations.

### 7. H-EV isolations provide lower miRNA diversity but may restore miRNA specificity

To evaluate the potential of NEXPLOR for biomarker discovery using additional omics layers, small RNA was isolated from our second cohort subset, comprising 8 HCC patients, 8 cirrhosis patients, and 4 healthy donors. Once again, samples underwent bulk EV isolation; one aliquot was kept for analysis and the other was further processed for h-EV isolation in a paired design. Samples were analyzed by small RNA-seq (**Figure 7A**).

**Figure 7.**
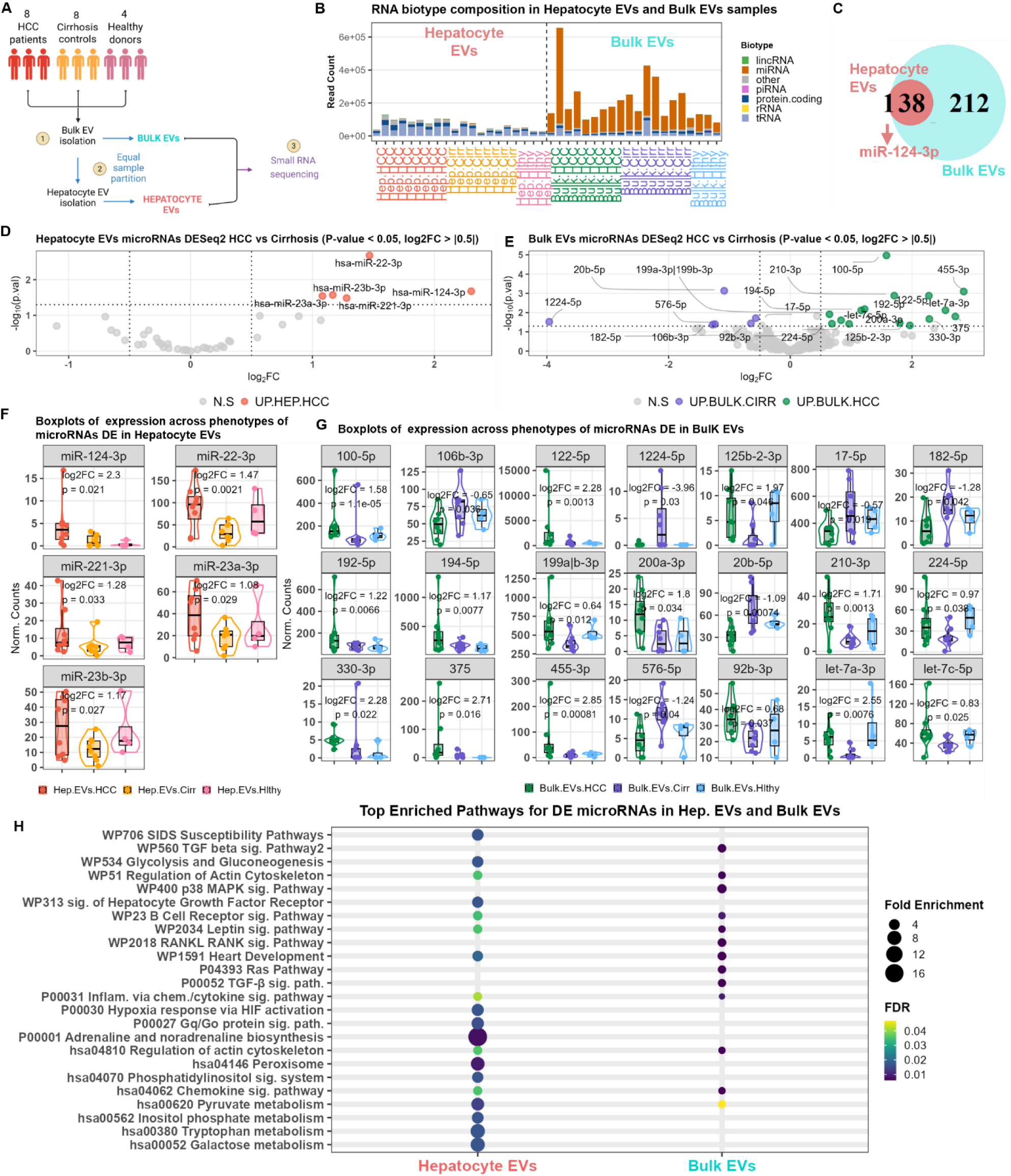
**Small RNA sequencing reveals differentially abundant miRNAs and distinct pathway enrichments between hepatocyte-enriched and bulk EVs in HCC versus cirrhosis**. **(A)** Schematic workflow for EV isolation and comparative small RNA sequencing (RNA-seq). Plasma samples were obtained from HCC patients (n = 8), cirrhosis controls (n = 8), and healthy donors (n = 4). EVs were first isolated as bulk EVs, then samples were equally partitioned, with hepatocyte EV isolation being applied to one portion. **(B)** Stacked barplot with RNA biotype composition of mapped reads on each sample. Hepatocyte EV samples correspond to the first 20 samples on the left and remaining correspond to bulk EV samples. X-axis displays read count with miRNA counts corresponding to the orange red portion of the bars. **(C)** Venn diagram displaying the overlap of miRNAs identifications (after filtering for low counts) between hepatocyte and bulk EV samples. Red: 1 miRNA (has-miR-124-3p) uniquely identified in hepatocyte EV samples; blue: 212 miRNAs uniquely identified in bulk EV samples; grey: 38 miRNAs identified in both. **(D)** Volcano plot of differentially abundant (DE) analysis for HCC vs cirrhosis in hepatocyte EV samples. miRNA significantly upregulated for HCC, when considering a p-value threshold of 0.05 and a log2 fold-change (log₂FC) of 0.5, are colored in red, non-significant are displayed in gray. **(E)** Volcano plot of DE analysis for HCC vs cirrhosis in bulk EV samples. miRNA significantly upregulated for HCC, when considering a p-value threshold of 0.05 and a log₂FC of 0.5, are colored in green, downregulated (upregulated in cirrhosis) are colored in purple, non-significant are displayed in gray. **(F)** Boxplot with levels of normalized counts (x-axis) across phenotypes, HCC (red), cirrhosis (orange) and healthy (pink) for the miRNAs found DE for HCC vs cirrhosis in hepatocyte EV samples. **(G)** Boxplot with levels of normalized counts (x-axis) across phenotypes, HCC (green), cirrhosis (purple) and healthy (light blue) for the miRNAs found DE for HCC vs cirrhosis in bulk EV samples. **(H)** Dot plot showing the top 12 significantly enriched Panther, Wiki and KEGG pathways (Benjamini-Hochberg adjusted p-value < 0.1,) associated with two sets of DE miRNAs for HCC vs cirrhosis samples, in hepatocyte and bulk EV samples. Dot color indicates false discovery rate (FDR), and size reflects fold enrichment.

After aligning reads and assessing the number of mapped reads it was clear there was a difference between the number of mapped reads for miRNA between bulk and h-EV samples, with a considerable high number of miRNA reads for bulk EVs, while the amount of reads for protein coding RNA and tRNA was similar across the two groups (**Figure 7B**). This is further demonstrated by the miRNA library size on **Figure S9A**.

After filtering out miRNA with low counts, 39 different miRNAs remained for hepatocyte EV samples while 250 remained for bulk EV samples, with 212 being found exclusively in bulk EVs. Interestingly, the unique miRNA found exclusively in h-EVs, *hsa-miR-124-3p* has been implicated as a tumor suppressor in HCC ^73^(**Figure 7C**).

Similar functional enrichment profiles were found for the targets of the 39 miRNAs found in h-EV samples and the 212 miRNAs exclusive to bulk EVs. (**Figure S9B and S9C**).

Differential abundance with DESeq2 ^74^ revealed 5 miRNAs, *hsa-miR-22-3p*, *hsa-miR-23b-3p*, *hsa-miR-124-3p*, *hsa-miR-23a-3p* and *hsa-miR-221-3*, differentially expressed in HCC vs Cirrhosis exclusively for h-EV samples (**Figure 7D**). No DE is determined for bulk EVs when comparing the same HCC and Cirrhosis patient samples. Nonetheless, for bulk EVs, 6 downregulated and 15 upregulated miRNAs for HCC were found (**Figure 7E**). Boxplots displaying normalized counts across the three phenotype subgroups (HCC, cirrhosis, and healthy) for both h-EVs and bulk EVs illustrate the differences of each DE miRNA (**Figures 7F and 7G**, respectively). DE results details can be found in **Tables S9 and S10**.

To investigate the biological pathways potentially modulated by these miRNA s, enrichment analysis was performed using the miRNA enrichment analysis and annotation tool (miEAA) ^75^, which incorporates both predicted and experimentally supported targets. In the comparison between HCC and cirrhosis, miRNAs upregulated in h-EVs, were uniquely enriched for cancer- and liver-relevant signaling pathways, including glycolysis and gluconeogenesis (WP534), hypoxia response via HIF activation (P00031), hepatocyte growth factor receptor signaling ^76^, and the GG/GO signaling pathway^77,78^, all of which have been previously implicated in HCC progression. Notably, these enrichments were exclusive to the h-EVs and not observed in the bulk EV-derived miRNA set. In contrast, pyruvate metabolism was enriched in both hepatocyte and bulk EVs, suggesting some shared metabolic deregulation across EV compartments in liver disease (**Figure 7H**).. Additional metabolic pathways, including inositol phosphate metabolism, tryptophan metabolism, and galactose metabolism, were also exclusively enriched in h-EVs. Similarly, the phosphatidylinositol signaling pathway, which feeds into the PI3K/AKT/mTOR axis, is frequently altered in HCC and contributes to tumorigenesis by promoting cell growth, survival, and metabolic rewiring ^79,80^. In contrast, the MAPK and TGF beta signaling pathways were uniquely enriched in the bulk EV-derived miRNA profile, suggesting that bulk EVs may capture pathway activity from non-hepatocyte sources or reflect a broader cellular response to liver injury and inflammation. The exclusive detection of metabolic and signaling pathways such as HIF activation, Hepatocyte growth facto receptor signaling and phosphatidylinositol signaling in h-EVs underscores the specificity of cell-type–targeted EV isolation in uncovering HCC-associated molecular mechanisms that may be diluted or absent in analyses of heterogeneous bulk EV populations.

## DISCUSSION

Resolving circulating ‘liquid’ biomarkers as signals for tissue-specific states is one of the grand challenges in precision diagnostics and precision medicine. Circulating EVs are naturally tissue- and tumor-informative but non-invasive. EVs may present advantages over tumor-agnostic methods using cf/ctDNA as their tissue of origin can be resolved via tissue-enriched or specific biomarkers, facilitating their isolation and application. This study presents what we believe to be the most comprehensive validation to date of the targeted isolation of EVs from a specific human tissue (hepatocytes) via the NEXPLOR platform, validating that EVs not only hold potential but applicability in this era. This type of methodology is particularly appealing in early cancer detection and monitoring especially when direct tissue access is too invasive or impractical.

The analysis of ctDNA circulating in blood has recently prompted an update to the National Comprehensive Cancer Network (NCCN) clinical guidelines for colorectal cancer (CRC) screening (https://www.nccn.org/guidelines/guidelines-detail?category=2&id=1429). CRC is characterized by a CpG island methylator phenotype (CIMP), resulting in consistent hypermethylation of key genes, alongside frequent mutation in others. CRC is also one of the highest DNA shedders among all cancers.

Unlike CRC, many cancers are not CIMP defined, genetic alterations are not as frequent or stable and shed low amounts of ctDNA, especially at early stages. CRC screening via liquid biopsy still requires large blood volumes (30-80 mL), altogether making ctDNA tests seemingly more challenging for other cancers. Some examples of low DNA shedders are gliomas, renal cell carcinoma, NSCLC and HCC, as previously discussed ^6,14–16^

Biopsies are also not commonly performed in many of those previous examples, making genomic tumor-informed assays for monitoring and precision medicine unfeasible.

Several studies suggest that EVs of defined tissue origins, may carry molecular surrogates of tissue states in the form of proteins, nucleic acids including DNAs and RNAs, lipids, and metabolites, protected within lipid bilayers which preserve molecular integrity.

Other freely-circulating biomarkers, such as soluble proteins, mRNAs and miRNAs usually lack spatial and biological resolution, limiting their ability to map molecular changes to specific tissues. More recently, bodies of evidence have shown miRNAs are widely present in EV isolations, and the possibility of analyzing circulating miRNAs derived from a known tissue is appealing in diagnostics.

Despite the intrinsic advantages of EVs, interrogating molecular insights from circulating EVs of a defined human tissue has remained challenging.

Circulating EVs are dominated by hematopoietic and vascular sources, exceeding organ-derived EVs by up to 5 orders of magnitude. For this reason, the immunocapture of circulating EV sub-types from bulk EV preparations, i.e. tagged with markers that reveal tissue-of-origin has been a central pursuit in EV-based liquid biopsies.

In this study, we presented NEXPLOR, adapted for the capture and isolation of hepatocyte- EVs (h-EVs). This provides rigorous validation of the isolation of circulating EVs from a defined tissue and illustrates the potential of circulating tissue-EVs to unlock clinically meaningful insights. Specifically we identified multiple central liver biology pathways, and, in subgroup analysis, central HCC and cirrhosis pathways identified by biomarkers with applications in early cancer detection, companion diagnostics and monitoring. We also attributed specificity to miRNAs, which is so far, the bottleneck for their clinical translation. Our approach combined *in silico* target discovery (**Figure 2**) and the experimental validation of 23 h-EV specific target candidates (**Table 1**, **Figure 3A**) with O-NEXOS. Four markers emerged as the most h-EV specific, namely ASGR1, ASGR2, TFR2 and SLCO1B1.

Together, NEXPLOR beads targeting the four markers were assembled as a final TOP4 h- EV capture cocktail (**Figure 3B**). The individual targets and cocktail mixture were assessed for EV capture on EP preparations isolated from snap-frozen Liver (healthy and diseased) and non-Liver healthy tissues, more precisely Kidney, Lung, Breast, Adipose and Muscle Tissues (**Figure 3C**). These tissues were selected as biologically relevant comparators due to their diverse developmental origins, metabolic activity, and potential to release tissue-specific EVs into circulation, allowing assessment of the specificity and selectivity of hepatocyte-derived EV isolation ^81^.

At equivalent extracellular particle (EP) concentrations, normalized to 1×10¹⁰ and 4×10⁹ particles/mL across all studied tissues, EV isolation from liver tissues consistently resulted in substantially higher yields than from all non-hepatic tissues evaluated (**Figure 3D**), supporting the developed methodology. The combined TOP4 h-EV capture cocktail, consistently enriched hepatocyte-derived EVs across healthy liver, cirrhosis, and HCC states, demonstrating robustness across disease contexts.

Next, we assembled a clinical cohort split into two patient subsets for hepatocyte-derived EV (h-EV) isolation from bulk EVs in plasma samples: one subset for proteomics analysis (**Figures 4A**), and the other for transcriptomic focused on miRNAs (**Figure 7A**). The aim was to compare the depth of molecular signals originating in the liver compared to the analysis on bulk EVs, for assessing liquid biopsy applications. The cohorts consisted of HCC patients (with a cirrhotic background), Cirrhotic patients (without HCC) and healthy individuals. HCC is a disease of hepatocytes, which constitute ∼ 80 % of the liver mass, 90 % of primary liver tumors are HCC and 85 % of HCC patients have underlying cirrhosis, the major risk factor for HCC development. ^82,83^

Novel biomarkers for the early detection and monitoring of HCC are urgently needed ^84^ We reasoned that our strategy for h-EV isolation could facilitate the identification of disease precision markers, manifesting the tissue state.

In fact, by capturing h-EVs, we obtained almost twice as many protein identifications compared to the bulk EV isolation, using the same starting samples, via TimsTOF SCP (**Figure 4B**). H-EV isolations also enabled clearer separation between clinical phenotypes (**Figure 4C**) and lower-intragroup variability (**Figure 4D**). GO-term analysis of proteins enriched or depleted on h-EV isolations, compared to bulk EVs, showed a depletion in plasma contaminant signals and increased enrichment of EV- and EP-related terms (**Figure 4F**). These results were further validated using the MISEV2023-compliant EV characterization framework (**Table S5**) ^8^ (In our datasets, 33 protein IDs categorized as 1a,1b,1c EV membrane markers and 2a,2b EV cytosolic markers, were found. Out of the 33 proteins, 18 were differentially elevated in h-EVs (p-value < 0.05), including CD81, CD82, LAMP1 and LAMP2 ^85,86^ 55 protein IDs in our datasets, categorized as 3a, 3b, 3c non-EV aggregates and 5a, 5b, 5c Secreted proteins were found, 23 of which were depleted from h-EV isolations (**Figure 5A**).

Highly expressed Hepatocyte proteins with cytosolic and transmembrane subcellular location were also differentially elevated on h-EV isolations (**Table S5**), such as GGH (transmembrane), TFR2 (transmembrane), ARG1 (cytosol), CES1 (transmembrane) or CAT (vesicles) and differentially decreased on secretory proteins like serine protease inhibitors and coagulation factors (**Figure 5B**). In all, enriching for h-EVs, resulted in an association with multiple metabolic KEGG pathways, consistent with the liver’s central metabolic role (**Figure 5C**) ^87,88^.

We applied the same comparative pipeline to stratify HCC and cirrhosis patient groups. In both, more unique protein groups (PGs) were identified in h-EV isolation than on bulk EVs (**Figure 6A**). In HCC, GO-term analysis of uniquely found PGs, strongly suggested association to “Extracellular exosome”, “Vesicle” and “Liver” terms, while unique PGs in bulk EVs of HCC found none (**Figure 6B, 6C**). Of note, “RNA binding”, “nucleic acid binding” terms were also strongly correlated to the HCC unique PGs, such as SYNCRIP and hnRNP family members, particularly HNRNPA2B1, supporting findings on selective miRNA sorting^89–91^_._

Mitochondrial-derived EV markers VDAC1 and VDAC2^92^ were almost exclusively detected in h-EV isolates, suggesting the capture of oxidative stress and potential metabolic dysfunction states through h-EVs (**Table S7**).

KEGG pathway analysis of disease-stratified comparisons reinforced these findings. Ferroptosis, a pathway central to fibrotic progression ^93^ was exclusively enriched in h-EVs of cirrhotic patients. “Central carbon metabolism in cancer”, a hallmark of liver cancer progression ^70,71^ was exclusively found in h-EVs of HCC patients. HIF-1 signaling pathway was only identified in h-EV isolations, being it enriched in HCC compared to Cirrhotic patients, which is consistent with HCC progression (**Figure 6F**)^94^.

Our cohort subset for miRNA investigations was relatively smaller (**Figure 7A**) and a considerable higher number of mapped miRNA reads was found for bulk EVs, with only 39 miRNAs being consistently represented in h-EV isolations. From the 39 miRNAs, 5 were significantly upregulated in h-EVs from HCC patients compared to Cirrhotic and healthy control group h-EVs (**Figure 7D, 7F**). These miRNAs were not DE between groups on bulk EVs (**Figure 7E**). The set included miR-124-3p and miR-23b-3p, both of which have been previously described as tumor suppressors in HCC but cellular disposal via exosomes has been linked to the acquisition of metastatic properties ^95,96^.

Differential Abundance in EVs from a diseased tissue, which otherwise their presence should suggest tumor suppressing mechanisms, may be a cancer-associated exporting mechanisms to decrease the cellular abundance and effect of these miRNAs in the tissue^97,98^.

Although h-EVs contained far fewer miRNAs compared to bulk EVs (**Figure 7B**), they were more informative by pathway enrichment analysis (**Figure 7H**).

Target enrichment analysis revealed a strong signal for pathways classically associated with HCC, including glycolysis and gluconeogenesis (WP534), hypoxia response via HIF activation (P00031), hepatocyte growth factor receptor signaling, and the GG/GO signaling pathway ^99–101^. Additionally metabolic pathways, such as inositol phosphate metabolism, tryptophan metabolism, galactose metabolism, phosphatidylinositol signaling pathway, central to the PI3K/AKT/mTOR axis ^102,103^ were also exclusively enriched in hepatocyte-derived EVs **(Figure 7H).** Illustrating that isolating a small, well-defined subpopulation of EVs, though low in miRNA content, can yield high-value molecular information when selectively captured.

It was not possible to measure the quantity of non h-EVs depleted from h-EV isolations but the body of evidence here presented strongly make a case that h-EV enrichment was indeed achieved enabling to zoom into liver tissue states, with for example, h-EVs from HCC patients showing HCC hallmark pathways and cirrhosis patients showing central cirrhosis pathways, strongly suggesting that well-validated tissue-EV enrichment platform enables the selective enrichment of tissue-specific signals across different omics (proteins and miRNAs).

Importantly in this work, NEXPLOR restores tissue specificity to circulating miRNAs, which are otherwise highly non-specific in plasma not only due to hematopoietic contributions but also because of overlapping miRNA circuitry across tissues^104^.

In conclusion, through organ-specific targeting, NEXPLOR enables DE analyses that more accurately reflect hepatocyte-driven biology across patient groups paving the way for early liver disease diagnosis, non-invasive deep-tissue proteomics and studying the involvement of miRNAs in liver diseases. This biological filtering may address a growing structural problem in AI-powered biomarker discovery pipelines: the instability and poor generalizability of models trained on weak, bulk-derived input layers lacking both functional and spatial resolution.

## MATERIALS AND METHODS

### Experimental Model and Human Samples Cell lines

Human cancer cell lines, MCF-7 (breast adenocarcinoma, HTB-22), BT-474 (breast ductal carcinoma, HTB-20), and HepG2 (hepatocellular carcinoma, HB-8065) were cultured in Advanced DMEM supplemented with 5% FBS and 4 mM GlutaMAX at 37°C in 5% CO₂.

### Tissue and plasma samples

Flash-frozen tumor and post-mortem tissues were sourced from commercial biobanks (TebuBio, Proteogenex, BioIVT, Amsbio, LifeNet). Plasma samples were collected from healthy donors and patients with cirrhosis or HCC via institutions including BIOBANCO, UCL Biobank, Imperial College London, and Novara University Hospital. All samples were collected with informed consent and processed per UK General Data Protection Regulation UK-GDPR guidelines.

### Extracellular vesicle isolation

#### From cell lines

At 70% confluency, cells were washed with PBS and cultured in serum-free medium. After 48 hours, the conditioned medium underwent sequential centrifugation (300 × g, 5 min; 3,000 × g, 10 min), filtration (0.22 µm), and concentration (Amicon Ultra-15 100K). EVs were isolated using Izon qEV SEC columns (70 nm for MCF-7/BT-474; 35 nm for HepG2) on an automatic fraction collector (AFC), eluted fractions pooled (1.5 mL), and stored at -80°C.

#### From tissues

Following digestion in Collagenase D (2 mg/mL) and DNase I (40 U/mL), samples were filtered (70 µm), washed, and centrifuged (500–10,000 × g). The supernatant was filtered (0.22 µm), concentrated, and processed with Izon qEV1 GEN2 35 nm columns to isolate EVs. Pooled fractions (2.8 mL) were stored at -80°C.

### From plasma

Thawed plasma was centrifuged (10,000 × g, 20 min), filtered (0.22 µm), and subjected to dual qEV 35 nm SEC runs. Fractions (5.6 mL total) were pooled and stored at -80°C.

### EV characterization

#### Nanoparticle tracking analysis (NTA)

Particle concentration and size distribution were determined using NanoSight NS300. Samples were diluted in PBS, measured in triplicate (3 × 60 s), and analyzed using NTA software v3.4 with a detection threshold of 7.

#### Western blotting

EVs (1 × 10¹⁰ particles/mL) were lysed in RIPA buffer, sonicated, and denatured. Proteins were resolved by SDS-PAGE, transferred to PVDF membranes, and probed using antibodies against CD9, CD63, and CD81. Chemiluminescence was detected using SuperSignal Pico substrate and imaged with G:BOX Chemi XRQ.

#### Bead preparation

Dynabeads® M-270 Epoxy MBs were conjugated with 10 µg antibody/mg beads using the Antibody Coupling Kit. Antibodies targeted EV surface markers and hepatocyte-specific proteins (e.g., ASGR1/2, SLCO1B1, TFR2). Beads were blocked and washed using buffers supplemented with 0.05% Tween®20.

#### TOP4 capture antibody cocktail

Antibodies against ASGR1, ASGR2, TFR2, and SLCO1B1 were conjugated separately and pooled in a 4:2:2:2 ratio to form the TOP4 hepatocyte EV capture cocktail.

### CD9^+^ EV isolation platforms

#### Mursla NEXPLOR workflow

EVs (7.9 × 10⁹ particles/mL) were incubated overnight with CD9-coupled MBs. Beads were washed and CD9^+^ EVs eluted with 0.2M glycine-HCl pH 2.5, neutralized with 1M Tris-HCl pH 8.0.

#### System Biosciences ExoFlow kit

Streptavidin MBs coupled with biotinylated anti-CD9 antibodies were used per SBI instructions. EVs were incubated overnight and eluted at 25°C using the kit’s buffer.

#### O-NEXOS detection of EV surface proteins

CD9 and CD63 were detected using biotinylated antibodies and magnetic bead capture. Incubations were performed in 96-well plates, followed by Poly-HRP strep addition and TMB-based colorimetric detection. Absorbance was measured at 450 nm using a SpectroStar Nano reader.

#### Liver-specific surface marker identification

A bioinformatic pipeline using Human Protein Atlas v20.1 identified liver-specific membrane proteins by combining mRNA expression specificity, IHC profiles, and UniProt membrane annotations. This yielded 81 candidate markers.

Expression validation was performed using Human Proteome Map and ProteomicsDB LC-MS/MS datasets. Proteins were ranked by liver specificity and visualized with log₂ intensity heatmaps.

### Proteomics and RNA-seq of h-EVs

#### Sample processing

SEC-isolated plasma EVs from HCC patients (n=13), cirrhosis patients (n=10), and healthy donors (n=6) were captured using TOP4 MBs. Lysates were prepared for MS and RNA-seq using RIPA buffer and processed via KingFisher Apex.

#### Mass spectrometry

Samples were processed with trypsin digestion and analyzed via data-independent acquisition (DIA) on a Bruker timsTOF-SCP. DIA-NN software (v1.8.1) was used for spectral library generation, identification, and quantification. MaxLFQ intensities were log₂-transformed.

#### Protein data analysis

IgG contaminants were removed, and PGs with <55% presence were excluded. Missing values were imputed (zero or left-censored). PCA, clustering, and Wilcoxon tests identified DE PGs (|fold-change| > 1, adj. p < 0.05). Functional analysis was conducted using clusterProfiler, FunRich, and STRING.

#### Hepatocyte-specific marker categorization

Differentially abundant PGs were categorized based on HPA data: hepatocyte-specific expression, secretion profiles, or both. Volcano plots and heatmaps visualized enrichment and annotation.

#### EV content characterization

Markers from MISEV2023 were mapped and visualized using EVqualityMS. Subtype distributions were analyzed by protein abundance.

### RNA isolation and sequencing

#### RNA extraction

RNA from NEXPLOR-captured EVs was extracted using IZON’s RNA Kit. SEC EVs underwent RNA extraction directly per kit protocol. RNA was eluted, dried, and resuspended for library prep.

#### Library preparation and sequencing

Small RNA libraries were generated using NEXTFLEX V4 with tRNA/YRNA blocking. Libraries were quantified and sequenced on the Illumina NextSeq 2000 platform to yield 50 bp single-end reads.

### Data analysis

Adapters were trimmed with BBDuk. Reads were aligned using exceRpt. miRNAs with low counts were filtered. DESeq2 was used for normalization and DE (|log₂FC| > 0.5, p < 0.05). Enrichment of predicted targets was performed with multiMiR and MiEAA using BH-corrected p-values.

### Statistical analysis

Statistical testing in **Figure 3C** and **Figure 3D** was performed using Kruskal-Wallis tests via kruskal.test and stat_compare_means from the ggpubr package (v0.6.0). Significance was set at p < 0.05.

## Supporting information

Table S1 - List of the database filtered 81 liver specific proteins (Rel. Fig 2)

Table S2 - Antibody validation table (Rel. Fig 3)

Table S5- Differentially abundance Bulk vs Hep. EVs results, MISEV2023 and hepatocyte expression annotations (Rel. Fig. 4, 5 and 6)

Table S6- KEGG pathways associated with protein-groups enriched in bulk and hepatocyte EV samples (Rel. Fig. 5)

Table S7 - Unique Protein Groups across liver disease groups (Rel. Fig. 6).xlsx

Table S8 - KEGG pathway associated with protein-groups enriched in bulk and hep. EV across liver disease phenotypes (Rel. Fig. 6)

Table S9- DESeq2 MicroRNAs Hepatocyte EVs HCC vs Hepatocyte EVs Cirrhosis results (Rel. Fig. 7)

Table S10- DESeq2 MicroRNAs Bulk EVs HCC vs Bulk EVs Cirrhosis results (Rel. Fig. 7)

## AKNOWLEDGMENTS

The authors would like to thank Dr. Ursula Arndt from the Illumina Ventures Lab (fka Illumina Accelerator) for facilitating access to their sequencing facilities and performing the sequencing of the samples in this study. The authors would also like to thank Evotec International GmbH for expert support with mass-spectrometry measurements, data acquisition, peptide identification, and protein quantification; Dr David James Pinato from Imperial College London and the GIMM Foundation Biobank in Lisbon for providing patient samples; and Tatiana Abdulkhalek at Mursla Bio for cross-checking all figure and table references and assisting with data assembly. Figures 1A, 1C, 1D, 1F, 2A, and 3B were created with BioRender.com.

