## Supplemental Figures S1-S9 for "Organ-specific isolation of hepatocyte extracellular vesicles from human plasma enables tissue-resolved proteomic and miRNA profiling"

Supplemental Information

### Targeted isolation of hepatocyte-derived extracellular vesicles from human plasma enhances the identification and discovery of extracellular vesicle, tissue and disease-specific proteins and miRNAs

Ricardo Figueiras[1], Susana Vagueiro[1], Raissa Kay[1], Elnaz Persia[1], Roger de Alwis[1], Razan Hijazi[1], Brian Davidson[2], Raquel Sanches-Kuiper[1], Pierre Arsène[1], Tomás Dias[1]

Figure S1 | Nanoparticle tracking analysis (NTA) of CD9$^{+}$ Hepatocyte extracellular vesicles (EVs).

Figure S2 | Western blot (WB) images of tetraspanins on EP preparations derived from cell lines.

Figure S3 | Expression of tetraspanin proteins ordered by tissue expression.

Figure S4 | Additional exploratory data analysis of proteomic results of liver disease and healthy plasma samples, processed with SEC or NEXPLOR.

Figure S5 | Hierarchical clustering heatmap of $log_2$-transformed intensity values for protein groups (PGs) categorized by MISEV2023 content-based criteria.

Figure S6 | Heatmap displaying scaled expression values (mass spectrometry) for bulk and hepatocyte EV samples, focusing on protein previously detected following immuno-isolation with the tetraspanins CD9, CD81, and CD63, as reported by Kowal et al. (2016).

Figure S7 | Hierarchical clustering heatmap of $log_2$-transformed intensity values for protein groups (PGs) with high protein expression and/or specific mRNA expression in hepatocytes, based on annotations from the Human Protein Atlas.

Figure S8 | STRING network and enrichment analysis of protein-groups (PGs) uniquely identified in hepatocyte or bulk EV samples from HCC or Cirrhosis patients.

Figure S9 | Small RNA sequencing based library sizes and functional enrichment of microRNAs after quality processing in liver disease and healthy plasma samples, processed with NEXPXLOR or SEC.

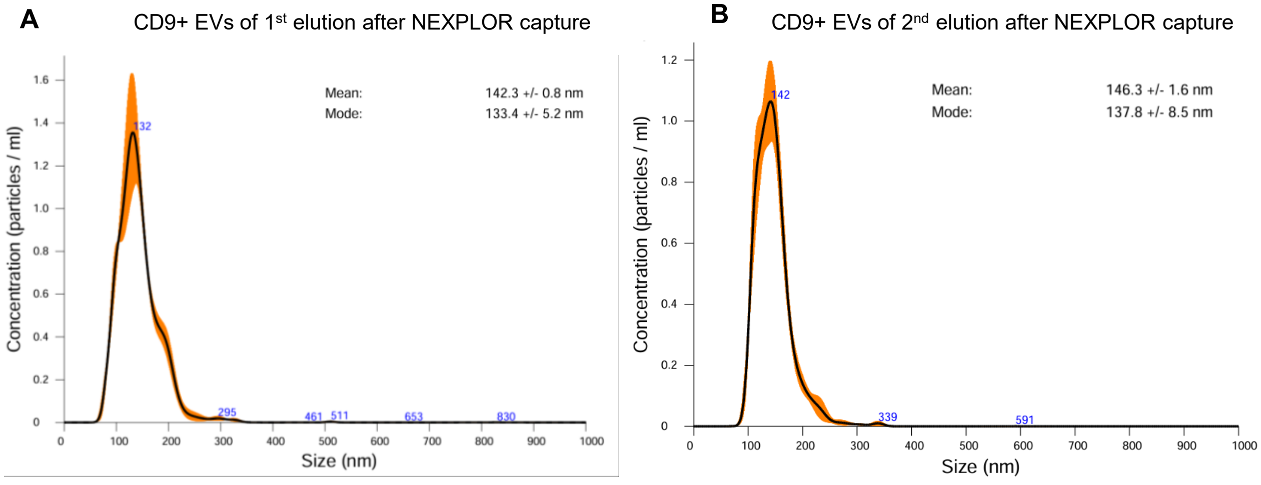


**Figure S1. Nanoparticle tracking analysis (NTA) of CD9$^{+}$ Hepatocyte extracellular vesicles (EVs). Related with Figure 1.**

NTA graphs showing particle size distribution (x-axis, in nanometers) and concentration (y-axis, in particles/mL) of CD9$^{+}$ hepatocyte EVs recovered after each elution step from magnetic beads using the NEXPLOR platform **(A)** Particle concentration and size profile following the first elution step **(B)** Particle concentration and size profile of EVs obtained through a second elution step, indicating additional vesicles released from beads after the first elution.

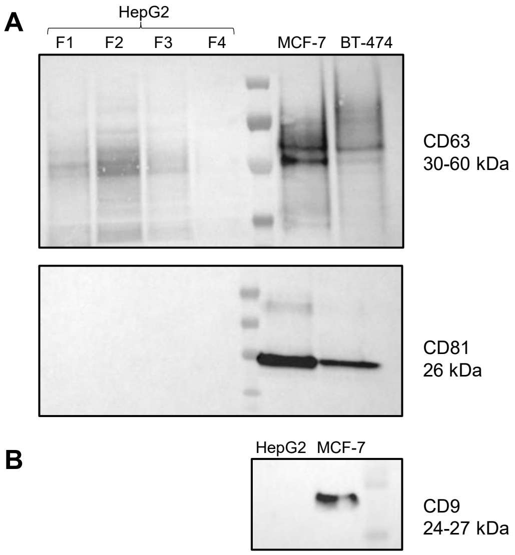


**Figure S2. Western blot (WB) images of tetraspanins on EP preparations derived from cell lines.**
**(A)** WB image for CD63 and CD81 protein. For HepG2, the EV fractions (F1, F2, F3) were analyzed separately as well as a non-EV fraction (F4). For MCF-7 and BT-474, the EVs were analysed only for the EV fractions already pooled **(B)** WB image for CD9 protein. The EV fractions were analyzed for HepG2 and MCF-7 already pooled.

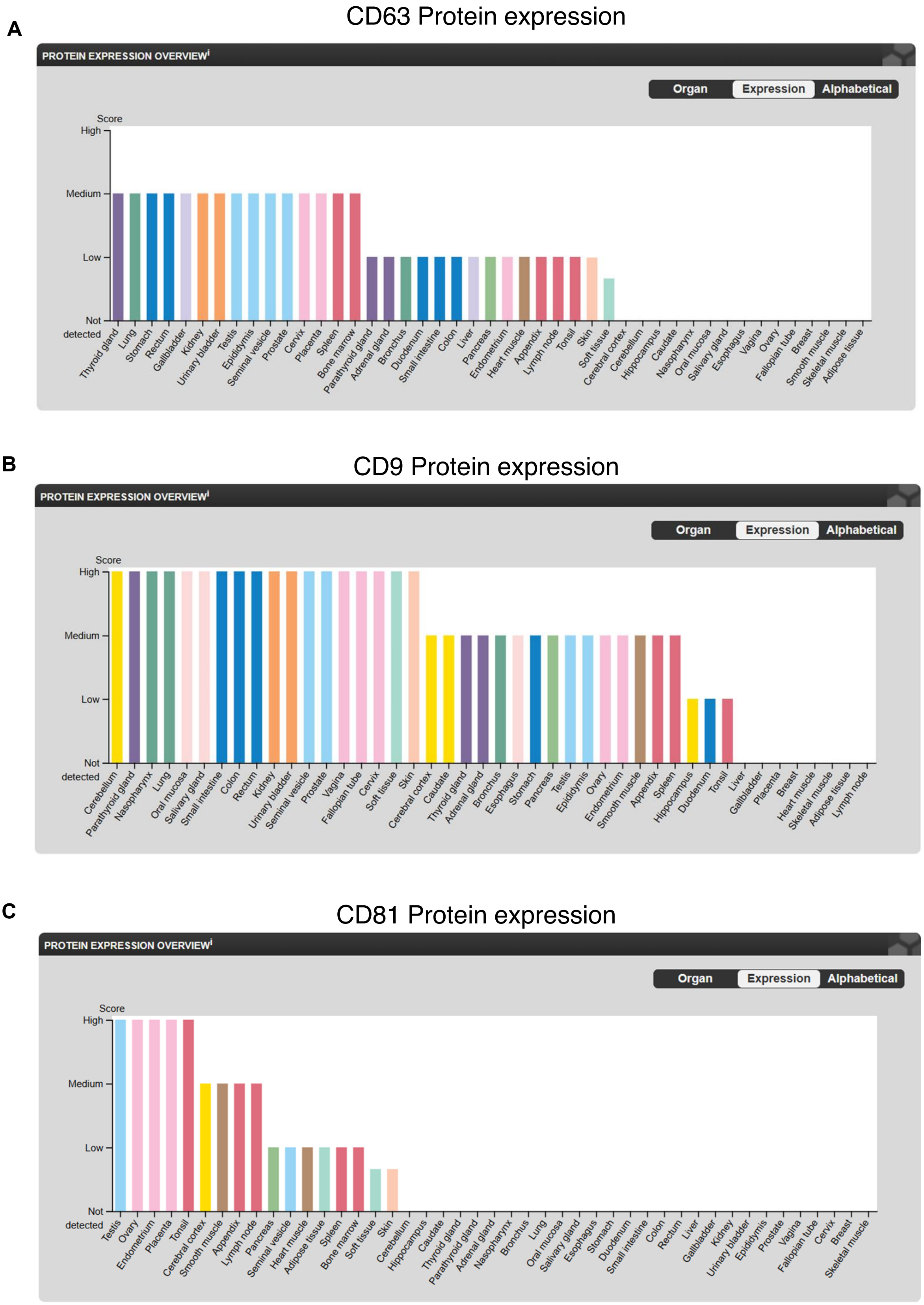


**Figure S3. Expression of tetraspanin proteins ordered by tissue expression.**
**(A)** Plot shows CD63 expression highlighting protein expression in the liver. Protein data was obtained from https://www.proteinatlas.org/ENSG00000135404-CD63/tissue **(B)** Plot shows CD9 expression highlighting absence of protein expression in the liver. Protein data was obtained from https://www.proteinatlas.org/ENSG00000010278-CD9/tissue **(C)** Plot shows CD81 expression highlighting absence of protein expression in the liver. Protein data was obtained from https://www.proteinatlas.org/ENSG00000110651-CD81/tissue. Data was last assessed in June 2025.

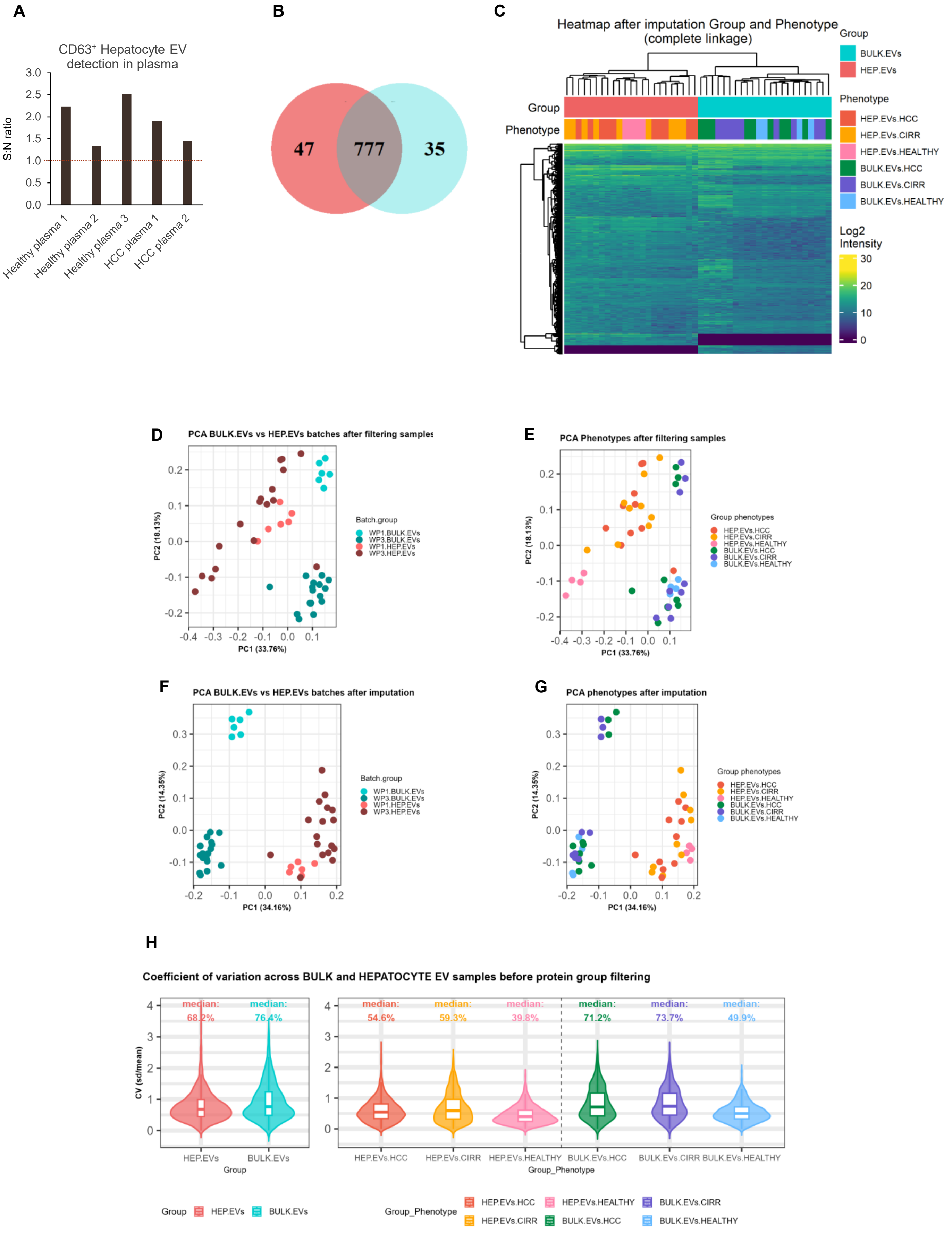
A
CD63⁺ Hepatocyte EV detection in plasma
S:N ratio
3.0
2.5
2.0
1.5
1.0
0.5
0.0
Healthy plasma 1
Healthy plasma 2
Healthy plasma 3
HCC plasma 1
HCC plasma 2
B
47
777
35
C
Heatmap after imputation Group and Phenotype (complete linkage)
Group
Phenotype
Group
BULK.EVs
HEP.EVs
Phenotype
HEP.EVs.HCC
HEP.EVs.CIRR
HEP.EVs.HEALTHY
BULK.EVs.HCC
BULK.EVs.CIRR
BULK.EVs.HEALTHY
Log2 Intensity
30
20
10
0
D
PCA BULK.EVs vs HEP.EVs batches after filtering samples
PC2 (18.13%)
PC1 (33.76%)
Batch.group
WP1.BULK.EVs
WP3.BULK.EVs
WP1.HEP.EVs
WP3.HEP.EVs
E
PCA Phenotypes after filtering samples
PC2 (18.13%)
PC1 (33.76%)
Group phenotypes
HEP.EVs.HCC
HEP.EVs.CIRR
HEP.EVs.HEALTHY
BULK.EVs.HCC
BULK.EVs.CIRR
BULK.EVs.HEALTHY
F
PCA BULK.EVs vs HEP.EVs batches after imputation
PC2 (14.35%)
PC1 (34.16%)
Batch.group
WP1.BULK.EVs
WP3.BULK.EVs
WP1.HEP.EVs
WP3.HEP.EVs
G
PCA phenotypes after imputation
PC2 (14.35%)
PC1 (34.16%)
Group phenotypes
HEP.EVs.HCC
HEP.EVs.CIRR
HEP.EVs.HEALTHY
BULK.EVs.HCC
BULK.EVs.CIRR
BULK.EVs.HEALTHY
H
Coefficient of variation across BULK and HEPATOCYTE EV samples before protein group filtering
CV (sd/mean)
median: 68.2%
median: 76.4%
median: 54.6%
median: 59.3%
median: 39.8%
median: 71.2%
median: 73.7%
median: 49.9%
HEP.EVs
BULK.EVs
Group
HEP.EVs.HCC
HEP.EVs.CIRR
HEP.EVs.HEALTHY
BULK.EVs.HCC
BULK.EVs.CIRR
BULK.EVs.HEALTHY
Group_Phenotype
Group
HEP.EVs
BULK.EVs
Group_Phenotype
HEP.EVs.HCC
HEP.EVs.HEALTHY
BULK.EVs.CIRR
HEP.EVs.CIRR
BULK.EVs.HCC
BULK.EVs.HEALTHY

**Figure S4. Additional exploratory data analysis of proteomic results of liver disease and healthy plasma samples, processed with SEC or NEXPLOR. Related to Figure 4.**
**(A)** Validation of hepatocyte EV capture and CD63+ EV detection using O-NEXOS. S:N values are calculated as CD63+ signal obtained after capture with the cocktail mix of the TOP4 antibody coupled magnetic beads divided by CD63+ signal obtained after capture with isotype antibodies **(B)** Venn diagram displaying the overlap of protein group (PG) identifications from mass spectrometry (MS) measurements of hepatocyte or bulk EV samples, following filtering for PGs present in ≥55% of the samples in either group. Red: PGs uniquely identified in hepatocyte EV samples; blue: PGs uniquely identified in bulk EV samples; grey: PGs identified in both **(C)** Hierarchical clustering heatmap of $log_2$-transformed intensity values for PGs (rows) and samples (columns), following filtering for PGs present in ≥55% of samples within at least one group and missing value imputation. Clustering was performed using Euclidean distance and complete linkage **(D)** Principal component analysis (PCA) of samples colored by MS batch (light red/blue – batch WP1; dark red/blue – batch WP3) and Group (red hues – hepatocyte EVs; blue hues – Bulk EVs) **(E)** PCA of samples colored by Group and Phenotype: red – hepatocyte EVs from HCC, “HEP.EVs.HCC”; orange – hepatocyte EVs from Cirrhotic liver, “HEP.EVs.CIRR”; pink – hepatocyte EVs from healthy liver, “HEP.EVs.HEALTHY”; green – Bulk EVs from HCC, “BULK.EVs.HCC”; purple – bullk EVs from cirrhotic liver, “BULK.EVs.CIRR”; light blue – Bulk EVs from healthy liver, “BULK.EVs.HEALTHY” **(F)** PCA of samples post-missing value imputation, colored as in (D) **(G)** PCA of samples post-missing value imputation, colored as in (E) **(H)** Violin plots and overlaid boxplots representing the coefficient of variation (CV) of PGs intensity values across samples, prior to PG filtering. Left panel: comparison of hepatocyte EVs (“HEP.EVS”, red) vs. bulk EVs (“BULK.EVs”, Blue). Right panel: stratification by group and phenotype (color scheme as in E/F). CV is shown on the Y-axis; median CV values are annotated as percentages above each group.

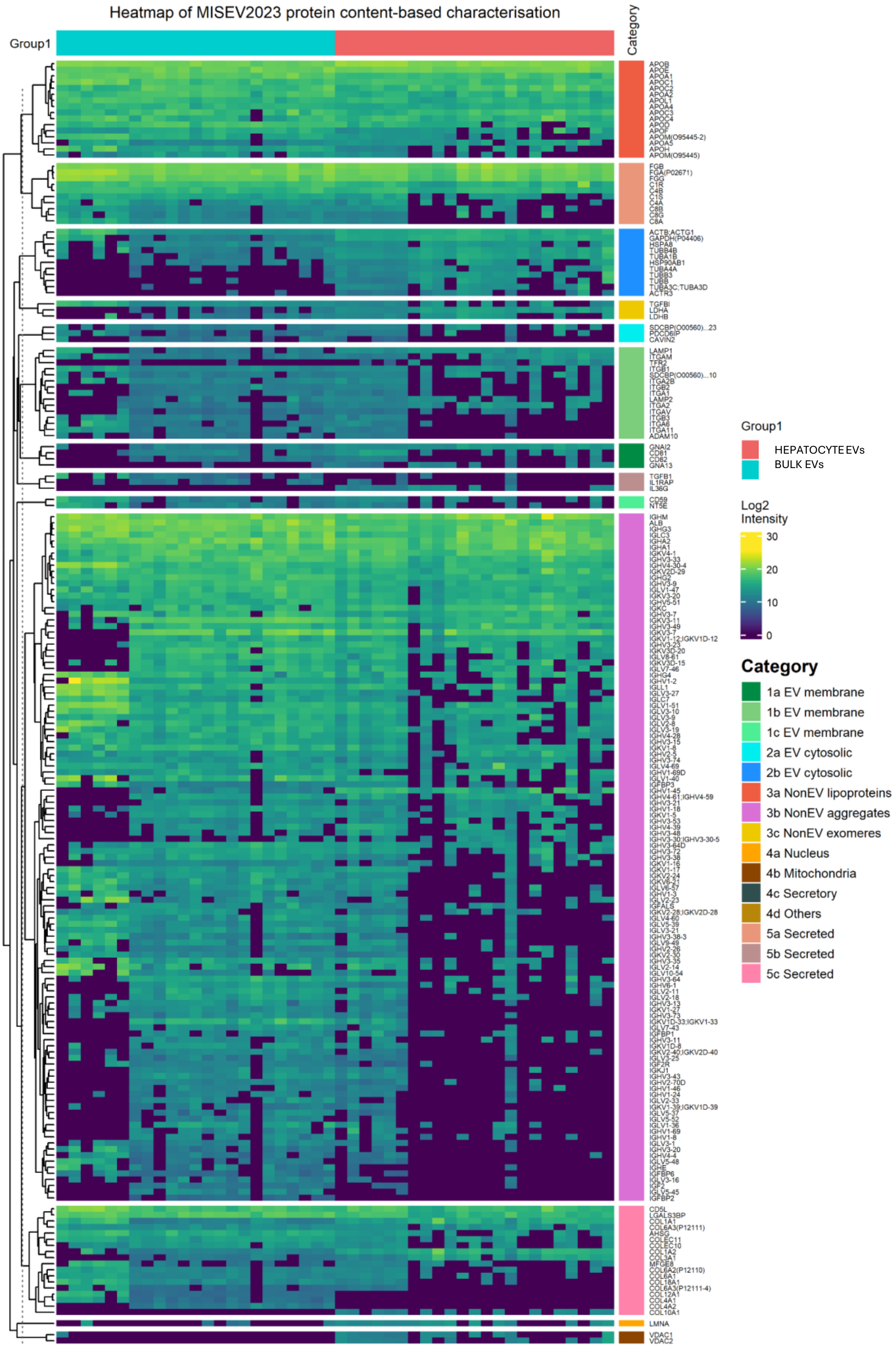

Heatmap of MISEV2023 protein content-based characterisation
Group1
Category
Group1
HEPATOCYTE EVs
BULK EVs
Log2 Intensity
30
20
10
0
Category
1a EV membrane
1b EV membrane
1c EV membrane
2a EV cytosolic
2b EV cytosolic
3a NonEV lipoproteins
3b NonEV aggregates
3c NonEV exomeres
4a Nucleus
4b Mitochondria
4c Secretory
4d Others
5a Secreted
5b Secreted
5c Secreted

**Figure S5. Hierarchical clustering heatmap of $log_2$-transformed intensity values for protein groups (PGs) categorized by MISEV2023 content-based criteria. Related to figure 5.**
Rows correspond to PGs (filtered for PGs present in ≥55% of samples in either group), annotated according to MISEV2023 extracellular vesicle (EV) content categories. PGs were clustered using Euclidean distance and complete linkage. Columns represent samples, color-coded by isolation method: red – Hepatocyte EVs ; blue – Bulk EVs. Rows categories are color-coded as follows: **Category 1**: Transmembrane (or GPI-anchored) proteins associated with plasma membrane and/or endosomes (**1a**: multi-pass transmembrane proteins, **1b**: single-pass transmembrane proteins, **1c**: GPI- or lipid-anchored proteins) – green hues; **Category 2**- Cytosolic proteins in EVs (**2a**: with lipid or membrane protein-binding ability, **2b**: promiscuous incorporation into EVs (and possibly NVEPs) – blue hues; **Category 3**- Major components of non-EV co-isolated structures (NVEPs) (**3a**: lipoproteins., **3b**: protein and protein/nucleic acid aggregates, **3c**: exomere or supermere-enriched components) – red, orchid, yellow; **Category 4**- Transmembrane, lipid-bound and soluble proteins associated with intracellular compartments other than PM/endosomes (**4a**: nucleus, **4b**: mitochondria, **4c**: secretory pathway, **4d**: others – orange, dark brown, dark grey, light brown. **Category 5**- Secreted proteins recovered with EVs (**5a**: blood-derived corona proteins, **5b**: cytokines and growth factors, **5c**: adhesion and extracellular matrix proteins) – pink hues.

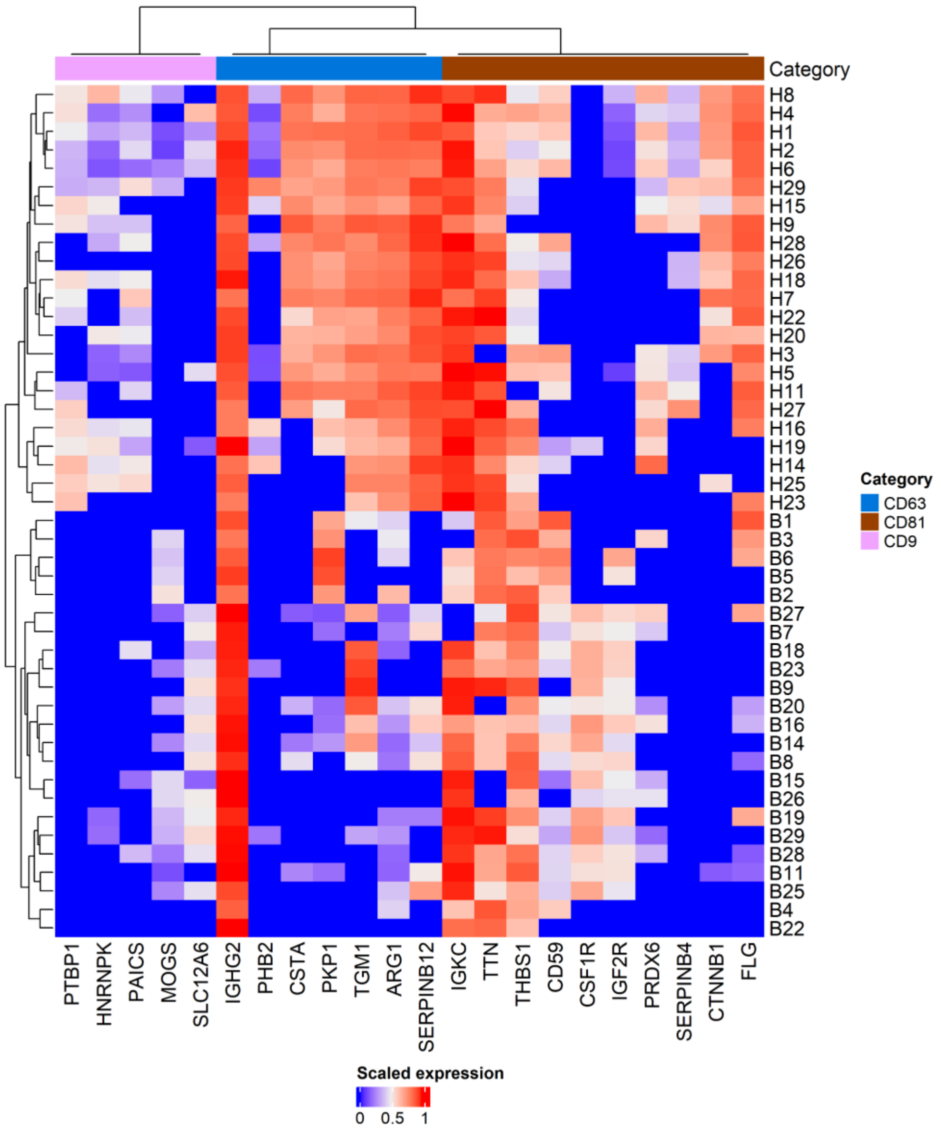


**Figure S6. Heatmap displaying scaled expression values (mass spectrometry) for bulk and hepatocyte EV samples, focusing on protein previously detected following immuno-isolation with the tetraspanins CD9, CD81, and CD63, as reported by Kowal et al. (2016). Related to Figure 5.**

Columns correspond to protein markers enriched through immuno-isolation with each respective tetraspanin. Column annotation indicate the targeted tetraspanin: blue – CD63, brown – CD81, pink – CD9.Rows correspond to sample IDs: H= Hepatocyte EVs samples, B= Bulk EV samples. Clustering was performed using Euclidean distance and complete linkage.

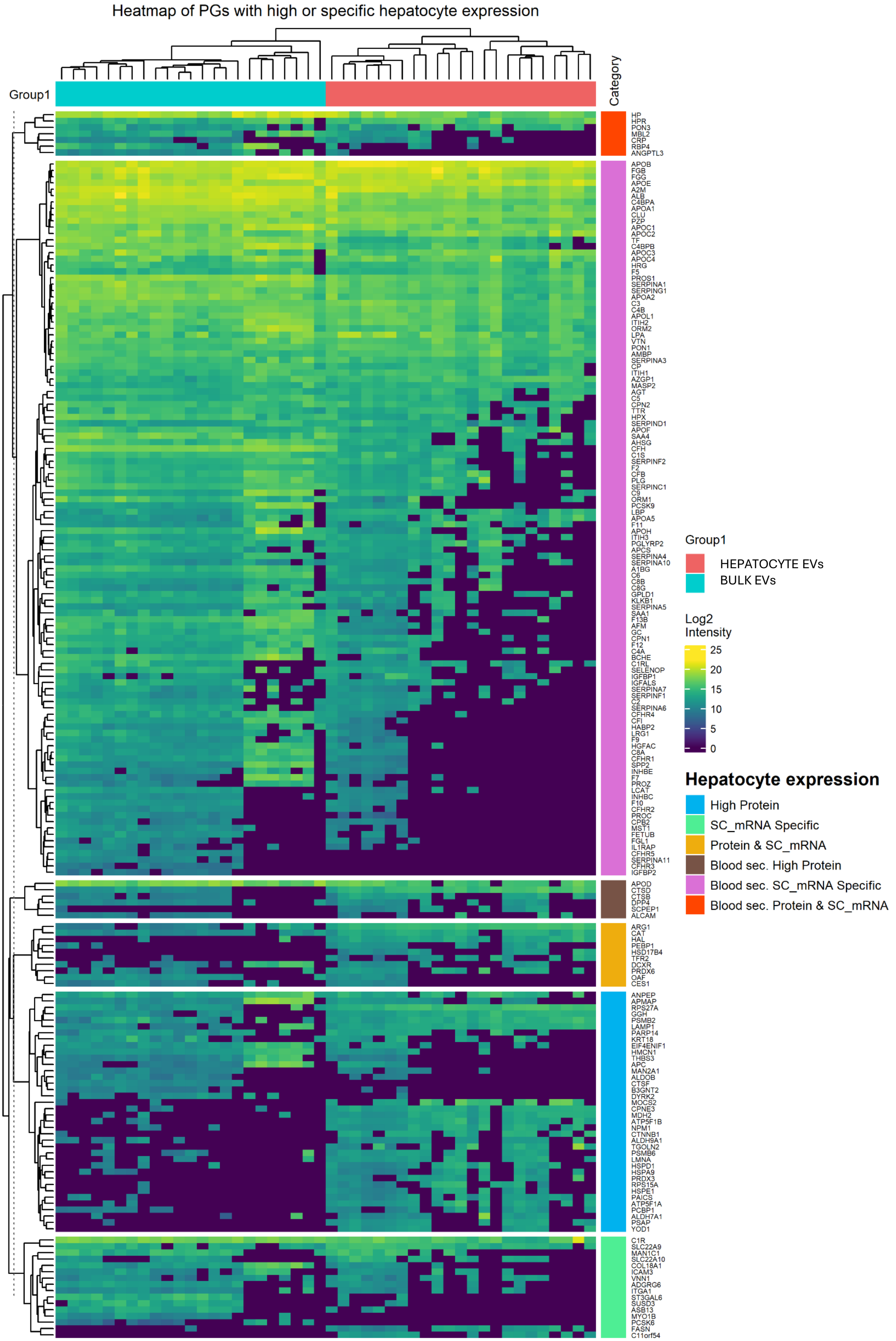
Heatmap of PGs with high or specific hepatocyte expression
Group1
Category
HP
HPR
PON3
MBL2
CRP
RBP4
ANGPTL3
APOB
FGB
FGG
APOE
A2M
ALB
C4BPA
APOA1
CLU
PZP
APOC1
APOC2
TF
C4BPB
APOC3
APOC4
HRG
F5
PROS1
SERPINA1
SERPING1
APOA2
C3
C4B
APOL1
ITIH2
ORM2
LPA
VTN
PON1
AMBP
SERPINA3
CP
ITIH1
AZGP1
MASP2
AGT
C5
CPN2
TTR
HPX
SERPIND1
APOF
SAA4
AHSG
CFH
C1S
SERPINF2
F2
CFB
PLG
SERPINC1
C9
ORM1
PCSK9
LBP
APOA5
F11
APOH
ITIH3
PGLYRP2
APCS
SERPINA4
SERPINA10
A1BG
C6
C8B
C8G
GPLD1
KLKB1
SERPINA5
SAA1
F13B
AFM
GC
CPN1
F12
C4A
BCHE
C1RL
SELENOP
IGFBP1
IGFALS
SERPINA7
SERPINF1
C2
SERPINA6
CFHR4
CFI
HABP2
LRG1
F9
HGFAC
C8A
CFHR1
SPP2
INHBE
F7
PROZ
LCAT
INHBC
F10
CFHR2
PROC
CPB2
MST1
FETUB
FGL1
IL1RAP
CFHR5
SERPINA11
CFHR3
IGFBP2
APOD
CTSD
CTSB
DPP4
SCPEP1
ALCAM
ARG1
CAT
HAL
PEBP1
HSD17B4
TFR2
DCXR
PRDX6
OAF
CES1
ANPEP
APMAP
RPS27A
GGH
PSMB2
LAMP1
PARP14
KRT18
EIF4ENIF1
HMCN1
THBS3
APC
MAN2A1
ALDOB
CTSF
B3GNT2
DYRK2
MOCS2
CPNE3
MDH2
ATP5F1B
NPM1
CTNNB1
ALDH9A1
TGOLN2
PSMB6
LMNA
HSPD1
HSPA9
PRDX3
RPS15A
HSPE1
PAICS
ATP5F1A
PCBP1
ALDH7A1
PSAP
YOD1
C1R
SLC22A9
MAN1C1
SLC22A10
COL18A1
ICAM3
VNN1
ADGRG6
ITGA1
ST3GAL6
SUSD3
ASB13
MYO1B
PCSK6
FASN
C11orf54
Group1
HEPATOCYTE EVs
BULK EVs
Log2
Intensity
25
20
15
10
5
0
Hepatocyte expression
High Protein
SC_mRNA Specific
Protein & SC_mRNA
Blood sec. High Protein
Blood sec. SC_mRNA Specific
Blood sec. Protein & SC_mRNA

**Figure S7. Hierarchical clustering heatmap of $\log_2$-transformed intensity values for protein groups (PGs) with high protein expression and/or specific mRNA expression in hepatocytes, based on annotations from the Human Protein Atlas. Related to figure 6.**

Rows correspond to PGs (filtered for PGs present in ≥55% of samples in either group), annotated based on hepatocyte expression features from the Human Protein Atlas. Columns represent samples, color-coded by sample type: red – Hepatocyte EVs; blue – Bulk EVs. Both PGs and samples were clustered by Euclidean distance and complete linkage.Rows categories are color-coded as follows: **High Protein** (light blue): High protein expression in hepatocytes. **SC_mRNA Specific** (light green): Hepatocyte-specific expression based on single-cell RNA sequencing. **Protein & SC_mRNA** (gold): Both high protein and single-cell RNA expression in hepatocytes. **Blood sec. High Protein** (dark brown): High hepatocyte protein expression, also secreted into blood. **Blood sec. SC_mRNA Specific** (purple): Hepatocyte-specific expression based on single-cell RNA sequencing and secreted into blood. **Blood sec. Protein & SC_mRNA** (red): Both high protein and single-cell RNA expression in hepatocytes whilst also known to be blood secreted.

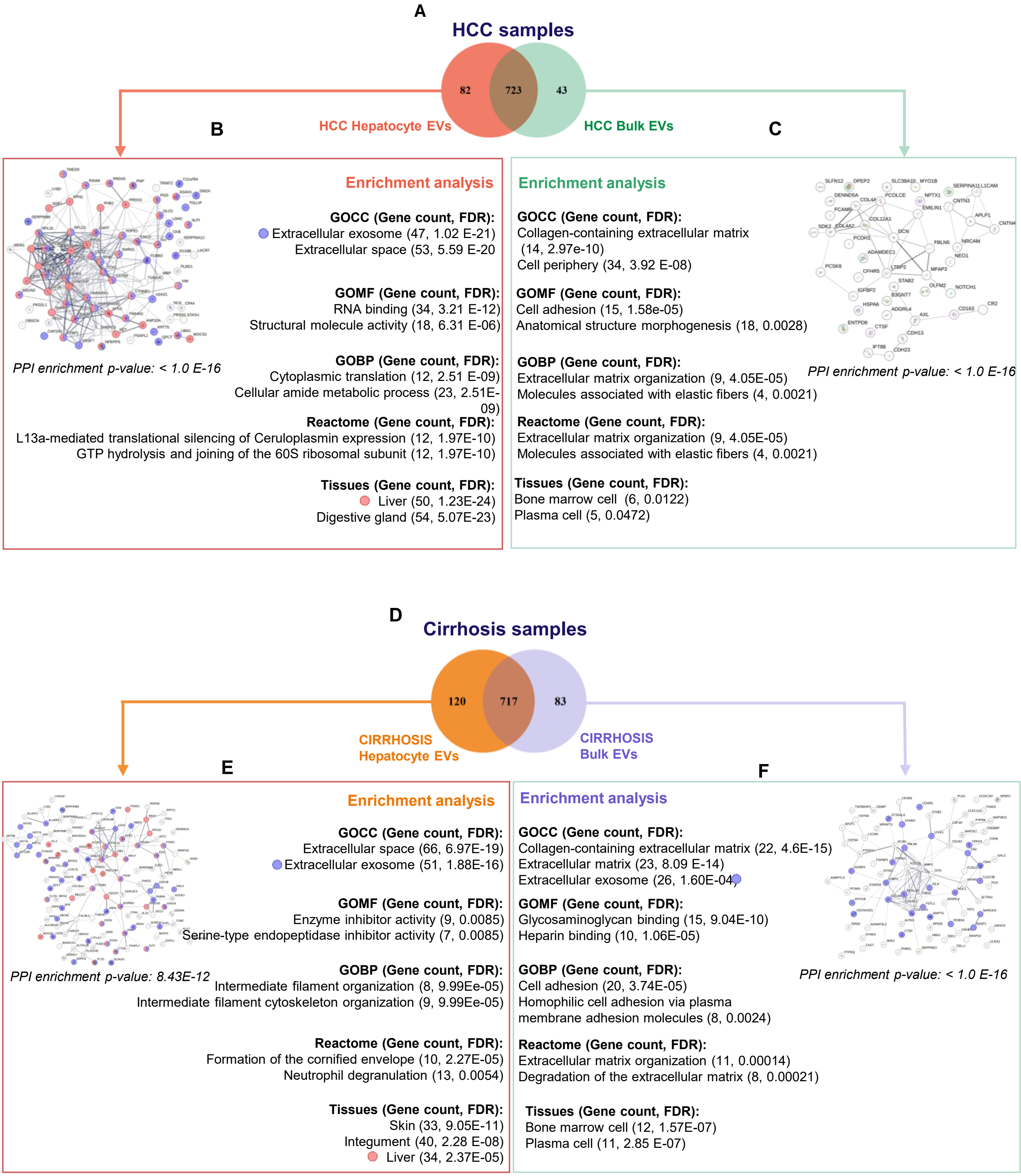

A
HCC samples
82
723
43
HCC Hepatocyte EVs
HCC Bulk EVs
B
C
Enrichment analysis
GOCC (Gene count, FDR):
Extracellular exosome (47, 1.02 E-21)
Extracellular space (53, 5.59 E-20
GOMF (Gene count, FDR):
RNA binding (34, 3.21 E-12)
Structural molecule activity (18, 6.31 E-06)
GOBP (Gene count, FDR):
Cytoplasmic translation (12, 2.51 E-09)
Cellular amide metabolic process (23, 2.51E-09)
Reactome (Gene count, FDR):
L13a-mediated translational silencing of Ceruloplasmin expression (12, 1.97E-10)
GTP hydrolysis and joining of the 60S ribosomal subunit (12, 1.97E-10)
Tissues (Gene count, FDR):
Liver (50, 1.23E-24)
Digestive gland (54, 5.07E-23)
PPI enrichment p-value: < 1.0 E-16
Enrichment analysis
GOCC (Gene count, FDR):
Collagen-containing extracellular matrix
(14, 2.97e-10)
Cell periphery (34, 3.92 E-08)
GOMF (Gene count, FDR):
Cell adhesion (15, 1.58e-05)
Anatomical structure morphogenesis (18, 0.0028)
GOBP (Gene count, FDR):
Extracellular matrix organization (9, 4.05E-05)
Molecules associated with elastic fibers (4, 0.0021)
Reactome (Gene count, FDR):
Extracellular matrix organization (9, 4.05E-05)
Molecules associated with elastic fibers (4, 0.0021)
Tissues (Gene count, FDR):
Bone marrow cell (6, 0.0122)
Plasma cell (5, 0.0472)
PPI enrichment p-value: < 1.0 E-16
D
Cirrhosis samples
120
717
83
CIRRHOSIS Hepatocyte EVs
CIRRHOSIS Bulk EVs
E
F
Enrichment analysis
GOCC (Gene count, FDR):
Extracellular space (66, 6.97E-19)
Extracellular exosome (51, 1.88E-16)
GOMF (Gene count, FDR):
Enzyme inhibitor activity (9, 0.0085)
Serine-type endopeptidase inhibitor activity (7, 0.0085)
GOBP (Gene count, FDR):
Intermediate filament organization (8, 9.99Ee-05)
Intermediate filament cytoskeleton organization (9, 9.99Ee-05)
Reactome (Gene count, FDR):
Formation of the cornified envelope (10, 2.27E-05)
Neutrophil degranulation (13, 0.0054)
Tissues (Gene count, FDR):
Skin (33, 9.05E-11)
Integument (40, 2.28 E-08)
Liver (34, 2.37E-05)
PPI enrichment p-value: 8.43E-12
Enrichment analysis
GOCC (Gene count, FDR):
Collagen-containing extracellular matrix (22, 4.6E-15)
Extracellular matrix (23, 8.09 E-14)
Extracellular exosome (26, 1.60E-04)
GOMF (Gene count, FDR):
Glycosaminoglycan binding (15, 9.04E-10)
Heparin binding (10, 1.06E-05)
GOBP (Gene count, FDR):
Cell adhesion (20, 3.74E-05)
Homophilic cell adhesion via plasma membrane adhesion molecules (8, 0.0024)
Reactome (Gene count, FDR):
Extracellular matrix organization (11, 0.00014)
Degradation of the extracellular matrix (8, 0.00021)
Tissues (Gene count, FDR):
Bone marrow cell (12, 1.57E-07)
Plasma cell (11, 2.85 E-07)
PPI enrichment p-value: < 1.0 E-16

**Figure S8. STRING network and enrichment analysis of protein-groups (PGs) uniquely identified in hepatocyte or bulk EV samples from HCC or Cirrhosis patients. Related to Figure 8.**
**(A)** Venn diagram displaying the overlap of protein group (PG) identifications from mass spectrometry (MS) measurements of hepatocyte or bulk EVs from HCC samples, following filtering for PGs present in ≥55% of the samples in either group. Red: PGs uniquely identified in HCC hepatocyte EVs; cyan: PGs uniquely identified in hepatocyte bulk EV samples; brown: PGs identified in both **(B)** STRING protein-protein interaction (PPI) network analysis (left) of the 82 PGs uniquely identified in HCC hepatocyte EVs compared to HCC bulk EVs. Enrichment analysis was performed using Gene Ontology cellular component (GOCC), molecular function (GOMF), biological process (GOBP), Reactome, and Jensen Tissues databases. The two top enriched terms for each category are shown (right). PGs annotated with the GOCC term “extracellular exosome” are colored in purple, and those associated with the Jensen Tissues term “Liver” are colored in red **(C)** STRING protein–protein interaction (PPI) network of 43 PGs uniquely identified in bulk EVs from HCC compared to hepatocyte EVs from HCC. Neither “extracellular exosome” (GOCC) nor “liver” (Jensen Tissues) terms were significantly enriched **(D)** STRING PPI network of 120 PGs uniquely identified in cirrhosis hepatocyte EVs compared to cirrhosis bulk EVs. Both “extracellular exosome” and “liver” terms were significantly enriched **(E)** STRING PPI network of 83 PGs uniquely identified in bulk EVs from cirrhosis compared to hepatocyte EVs from cirrhosis. The “extracellular exosome” term was significantly enriched, but “liver” was not.

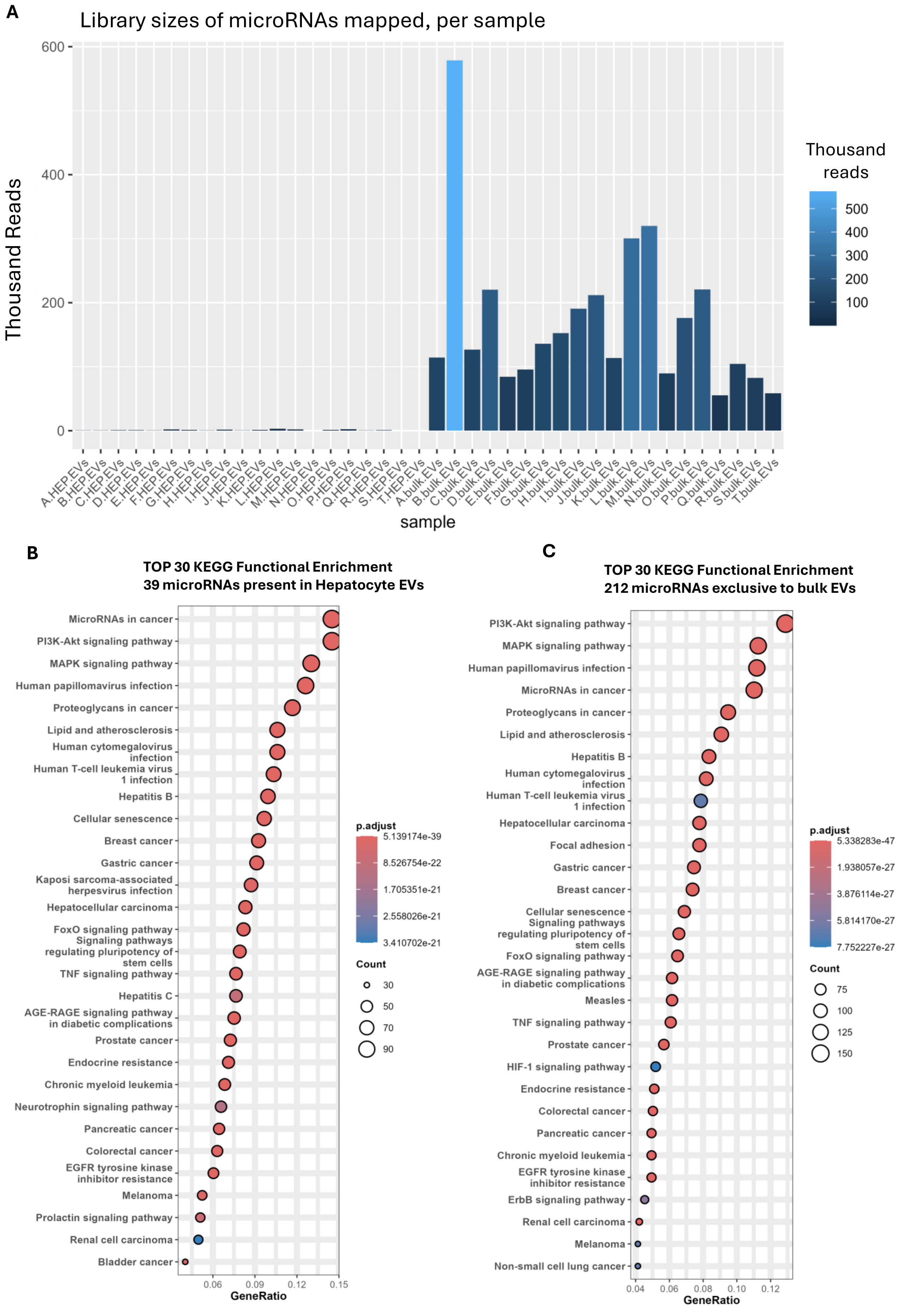

A
Library sizes of microRNAs mapped, per sample
Thousand Reads
600
400
200
0
Thousand reads
500
400
300
200
100
A.HEP.EVs B.HEP.EVs C.HEP.EVs D.HEP.EVs E.HEP.EVs F.HEP.EVs G.HEP.EVs H.HEP.EVs I.HEP.EVs J.HEP.EVs K.HEP.EVs L.HEP.EVs M.HEP.EVs N.HEP.EVs O.HEP.EVs P.HEP.EVs Q.HEP.EVs R.HEP.EVs S.HEP.EVs T.HEP.EVs A.bulk.EVs B.bulk.EVs C.bulk.EVs D.bulk.EVs E.bulk.EVs F.bulk.EVs G.bulk.EVs H.bulk.EVs I.bulk.EVs J.bulk.EVs K.bulk.EVs L.bulk.EVs M.bulk.EVs N.bulk.EVs O.bulk.EVs P.bulk.EVs Q.bulk.EVs R.bulk.EVs S.bulk.EVs T.bulk.EVs
sample
B
TOP 30 KEGG Functional Enrichment
39 microRNAs present in Hepatocyte EVs
MicroRNAs in cancer
PI3K-Akt signaling pathway
MAPK signaling pathway
Human papillomavirus infection
Proteoglycans in cancer
Lipid and atherosclerosis
Human cytomegalovirus infection
Human T-cell leukemia virus 1 infection
Hepatitis B
Cellular senescence
Breast cancer
Gastric cancer
Kaposi sarcoma-associated herpesvirus infection
Hepatocellular carcinoma
FoxO signaling pathway
Signaling pathways regulating pluripotency of stem cells
TNF signaling pathway
Hepatitis C
AGE-RAGE signaling pathway in diabetic complications
Prostate cancer
Endocrine resistance
Chronic myeloid leukemia
Neurotrophin signaling pathway
Pancreatic cancer
Colorectal cancer
EGFR tyrosine kinase inhibitor resistance
Melanoma
Prolactin signaling pathway
Renal cell carcinoma
Bladder cancer
0.06 0.09 0.12 0.15
GeneRatio
p.adjust
5.139174e-39
8.526754e-22
1.705351e-21
2.558026e-21
3.410702e-21
Count
30
50
70
90
C
TOP 30 KEGG Functional Enrichment
212 microRNAs exclusive to bulk EVs
PI3K-Akt signaling pathway
MAPK signaling pathway
Human papillomavirus infection
MicroRNAs in cancer
Proteoglycans in cancer
Lipid and atherosclerosis
Hepatitis B
Human cytomegalovirus infection
Human T-cell leukemia virus 1 infection
Hepatocellular carcinoma
Focal adhesion
Gastric cancer
Breast cancer
Cellular senescence
Signaling pathways regulating pluripotency of stem cells
FoxO signaling pathway
AGE-RAGE signaling pathway in diabetic complications
Measles
TNF signaling pathway
Prostate cancer
HIF-1 signaling pathway
Endocrine resistance
Colorectal cancer
Pancreatic cancer
Chronic myeloid leukemia
EGFR tyrosine kinase inhibitor resistance
ErbB signaling pathway
Renal cell carcinoma
Melanoma
Non-small cell lung cancer
0.04 0.06 0.08 0.10 0.12
GeneRatio
p.adjust
5.338283e-47
1.938057e-27
3.876114e-27
5.814170e-27
7.752227e-27
Count
75
100
125
150

**Figure S9. Small RNA sequencing based library sizes and functional enrichment of microRNAs after quality processing in liver disease and healthy plasma samples, processed with NEXPXLOR or SEC. Related to Figure 7.**

**(A)** Library size per sample, in thousands of reads, of microRNAs from miRNA sequencing mapped after pre-processing (adapter trimming, sequence alignment and filtering out low read counts). “HEP.EVs” are hepatocyte EVs enriched via NEXPLOR and “bulk.EVs” are bulk EVs enriched via SEC **(B)** Top 30 KEGG pathways enriched for by genes targeted through the 39 microRNAs found in HCC hepatocyte EVs **(C)** Top 30 KEGG pathways enriched for by genes targeted through the 221 exclusive microRNAs found in HCC bulk EVs
